# Optimizing progeny allocation strategies in breeding schemes while updating genomic prediction models

**DOI:** 10.1101/2025.08.19.671165

**Authors:** Kosuke Hamazaki, Koji Tsuda, Hiroyoshi Iwata

## Abstract

Genomic selection has revolutionized breeding by enabling the early identification of superior individuals using genome-wide markers, enhancing breeding efficiency and accelerating variety development. Over the past decade, new selection and mating strategies — leveraging optimization methods and other approaches — have been introduced to improve various decision-making processes in breeding programs. However, optimizing breeding remains challenging when the positions and effects of quantitative trait loci are unknown. We developed a framework that optimizes breeding strategies while updating genomic prediction models during breeding schemes. By implementing intermediate model updates, we enabled re-optimization of allocation strategies based on updated predictions. Our simulations compared this approach with equal allocation and optimal cross selection methods across various selection intensities and genetic architectures. Results demonstrated our optimized allocation strategy significantly outperformed the other approaches under moderate to low selection intensities, particularly when combined with model updates. While genetic gains plateaued without updates, our approach enabled continuous improvement through the final generation. The framework showed exceptional robustness across different simulation conditions and better maintained genetic diversity while controlling changes in population structure. This confirms that optimized allocation strategies remain effective when using estimated marker effects rather than true effects, providing a practical framework for improving real-world breeding programs.

## Introduction

As the global population grows, the demand for stable food supply increases, while climate change complicates food production (Godfray et al., 2010; Lobell et al., 2011; Wheeler and von Braun, 2013). This creates an urgent need to efficiently develop new crop cultivars with desired agricultural features, including environmental adaptability, high yield, and nutritional value (Tilman et al., 2011; Ray et al., 2013). Conventional breeding methods cannot address these challenges quickly enough, necessitating technological innovations (Tester and Langridge, 2010; Bailey-Serres et al., 2019; Varshney et al., 2021). In recent years, genomic selection (GS) has emerged as a breakthrough method that significantly enhances breeding efficiency (Meuwissen et al., 2001). GS utilizes whole-genome DNA marker information and previously trained genomic prediction (GP) models to predict genotypic values before phenotypic evaluation, enabling early selection of superior individuals (Meuwissen et al., 2001; Jannink et al., 2010; Crossa et al., 2017). This approach shortens breeding cycles, allowing for significant genetic gains in a short period (Heffner et al., 2009). GS has proven particularly valuable for improving complex quantitative traits such as environmental stress tolerance, yield, and nutritional value across numerous crop species (Bernardo and Yu, 2007; Heffner et al., 2011a; Onogi et al., 2015; Duhnen et al., 2017; Minamikawa et al., 2017; Yabe et al., 2018).

Since GS was introduced to crop breeding, numerous studies have aimed to improve selection efficiency through the enhancement of GP models (Meuwissen et al., 2001; Gianola and van Kaam, 2008; de los Campos et al., 2009; Jannink et al., 2010; Gianola et al., 2011; Habier et al., 2011; Heffner et al., 2011b). However, effectively implementing GS in breeding schemes spanning multiple generations requires not only better GP models but also well-designed selection and mating strategies based on GP results (Wang et al., 2018). For selection strategies, instead of simply using predicted breeding values by GP (GEBV) as in Meuwissen et al. (2001), methods such as WBV (Goddard, 2009; Jannink, 2010), OHV (Daetwyler et al., 2015), OPV (Goiffon et al., 2017), PCV (Han et al., 2017), and EMBV (Müller et al., 2018) have been devised that consider allele frequency and recombination to provide mid- to long-term advantages. Additionally, for mating strategies, several approaches have been proposed that directly identify superior crossing pairs by evaluating potential genetic variance in the progeny of crossed pairs through simulations (Lian et al., 2015; Mohammadi et al., 2015; Yao et al., 2018) or theoretical formulas (Zhong and Jannink, 2007; Lehermeier et al., 2017; Allier et al., 2019b).

In recent years, optimization methods for various decisions in breeding schemes have gained significant attention (Hamazaki and Iwata, 2021; Diot and Iwata, 2022; Moeinizade et al., 2022; Jannink et al., 2023; Hassanpour et al., 2025). For selection and mating strategies, optimal contribution selection, an early approach, optimizes the contribution of each genotype to the next generation by balancing genotypic gains with genetic diversity based on the pedigree information (Meuwissen, 1997). This method evolved to incorporate genomic information (Kemper et al., 2012) and developed into optimal cross selection (OCS), which selects ideal crossing pairs while balancing genotypic gains with genetic diversity (Kinghorn, 2011), with practical applications of OCS also being proposed (Gorjanc et al., 2018; Allier et al., 2019b, 2020). While research on mating design problem has a long history (Jansen and Wilton, 1985), the past decade has seen numerous new methods emerge alongside OCS, mainly driven by the utilization of genomic information and advancements in simulation technologies. Look-ahead selection predicts the genetic gains of new varieties produced several generations later under simplified assumptions to select optimal mating pairs (Moeinizade et al., 2019; Zhang and Wang, 2022). Cross potential selection combines estimated distribution of genotypic values for inbred progeny with integer programming to select multiple crossing pairs simultaneously under some constraints (Sakurai et al., 2024). Other approaches determine the optimal progeny allocations across mating pairs to achieve a Pareto front of expected genetic gain and variance in the next generation (Hunter and McClosky, 2016). Notably, Hamazaki and Iwata successfully integrated breeding simulations with black-box optimization to maximize the genetic gains of new varieties over multiple generations by optimally allocating progeny for each mating pair at each generation (Hamazaki and Iwata, 2024), and subsequently enhanced this approach by incorporating automatic differentiation, a state-of-the-art computational technology (Hamazaki et al., 2024).

However, many studies on decision-making in breeding programs often operate under the ideal assumption that true marker effects are known (Kemper et al., 2012; Daetwyler et al., 2015; Goiffon et al., 2017; Wang et al., 2018; Moeinizade et al., 2019, 2022; Amini et al., 2021; Zhang and Wang, 2022; Hamazaki and Iwata, 2024; Hamazaki et al., 2024). In other words, these studies propose breeding strategies assuming that the positions and effects of genes controlling traits can be completely understood. In reality, it is practically impossible to fully grasp the exact positions and effects of quantitative trait loci (QTLs) in advance, creating a significant disconnect between the above assumptions and the reality of actual breeding programs. Although fully identifying loci and effects for quantitative traits remains challenging even in major crops such as rice and maize, QTLs and genes have been identified for some traits and used in breeding through QTL pyramiding and marker-assisted selection (Collard and Mackill, 2008). For less-studied crops, such as neglected and underutilized species (NUS) (Padulosi et al., 2013), however, such discoveries are rare even for traits governed by just a few major genes, thereby broadening the range of traits where GP is particularly valuable. Therefore, evaluating breeding strategies based on marker effects estimated through GP has significant value, particularly for improving breeding efficiency in less-studied crops.

GP accuracy directly impacts selection efficiency, making it crucial when advancing breeding schemes that use marker effects estimated by GP. Generally, as breeding schemes progress and selection continues, alleles become fixed, and the genetic variance of the population decreases significantly due to the Bulmer effect (Bulmer, 1971; Van Grevenhof et al., 2012). This causes later generations to differ substantially from the initial population. When using a GP model trained only on the initial population to predict genotypic values of these later generations, prediction accuracy gradually declines with each generation. To maintain GP accuracy, it is necessary to update the GP model regularly throughout the breeding scheme. In fact, previous studies have demonstrated that these periodic model updates enable continuous genetic improvement, even in mid-term or long-term breeding schemes (Jannink, 2010; Gorjanc et al., 2018).

Based on this background, our study proposes a framework for optimizing breeding strategies while updating the GP model during the breeding scheme, assuming realistic situations where the QTL positions and effects remain unknown. Specifically, we focus on the approach developed by Hamazaki and Iwata (2024) for optimizing progeny allocation across mating pairs, and verify its effectiveness by re-optimizing the progeny allocation strategy in conjunction with model updates.

## Materials and Methods

Throughout the study, R version 4.4.1 (R Core Team, 2024) was used to simulate the marker genotype of the initial breeding population, implement breeding optimization, and conduct breeding simulations.

### System design of the study

As outlined by Hamazaki and Iwata (2024), breeders follow a breeding scheme based on the progeny allocation strategy proposed by an “AI breeder” (Supplemental Fig. 1). The AI breeder receives current breeding population data—including marker genotype data, phenotypic data of target traits, and recombination rates between genetic markers— from actual breeders and outputs a set of parameters representing the optimal allocation strategy. Below, we describe how the AI breeder optimizes the resource allocation of progeny.

After receiving the current population information at generation *t*, the AI breeder explores the optimal allocation strategy through repeated breeding simulations based on different allocation parameters, 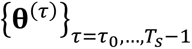 (Supplemental Fig. 1). The parameter vector **θ**^(*τ*)^ ∈ ℝ^*d*^ serves as the AI breeder’s decision-making framework. In these simulations, *τ* ∈ {*τ*_0_,…, *T*_*s*_ − 1} denotes the current generation number, where *τ*_0_ ∈ ℕ and *T*_*s*_ ∈ ℕ are the initial and final generation numbers. On the other hand, *t* ∈ {0,…, *T*_*r*_ − 1} denotes the current generation number in breeding schemes, which are conducted by actual breeders, where *T*_*r*_ ∈ ℕ is the final generation in actual breeding schemes.

In each breeding simulation, we assess the performance of allocation strategies by computing the genetic gain in the final generation *T*_*s*_ after multiple cycles of selection, mating, and allocation. The genetic gain, 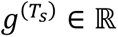, quantifies the improvement in genotypic values of the top 5 % of individuals in the final population compared to the initial population. Thus, each breeding simulation can be represented as a function with input 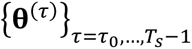 and output 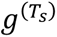, as in Equation 1.

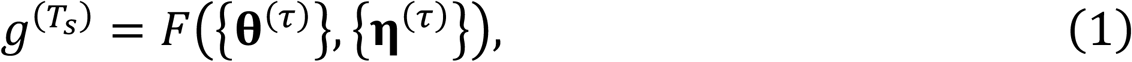

where *F* is a function representing each breeding simulation. Because breeding simulations incorporate random factors related to recombination events when generating new individuals from mating pairs in each generation *τ*, represented by random variables **η**^(*τ*)^ ∈ ℝ^*M*^ in this study. To evaluate the performance of {**θ**^(*τ*)^}, we must calculate the expectation with respect to {**η**^(*τ*)^} through repeated breeding simulations. Thus, the objective of the AI breeder is to optimize {**θ**^(*τ*)^} so that *F*({**θ**^(*τ*)^}) in the following Equation 2 is maximized.

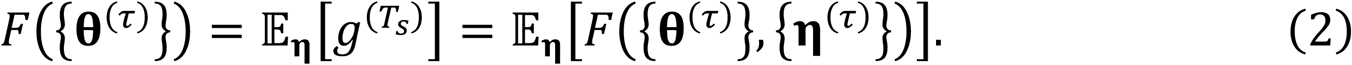

As noted by Hamazaki and Iwata (2024), this problem can be viewed as a black-box optimization problem where only the inputs and outputs of function *F* are observable. Therefore, we optimize the allocation strategy by combining repeated breeding simulations with StoSOO (Valko et al., 2013), a black-box optimization technique.

Here, since the GP model used in breeding simulations conducted by the AI breeder becomes less accurate over generations, it requires updates during the breeding scheme. Thus, in this study, we update the GP model at generation *t* = *t*_*u*_ (0 < *t*_*u*_ < *T*_*r*_), at which point the AI breeder re-optimizes the allocation strategy (Fig. 1). Specifically, the construction of the GP models and the optimization by the AI breeder occur twice— at generation *t* = 0 and *t* = *t*_*u*_. Throughout the study, we set *T*_*r*_ = 4 and *t*_*u*_ = 2. This approach allows us to examine how model updates affect the allocation strategy optimization by comparing results with and without model updates (Fig. 1 and Supplemental Fig. 2). We compared the optimized allocation strategy proposed by Hamazaki and Iwata (2024) with equal allocation and OCS under the breeding schemes with or without model updates.

**Fig. 1.**
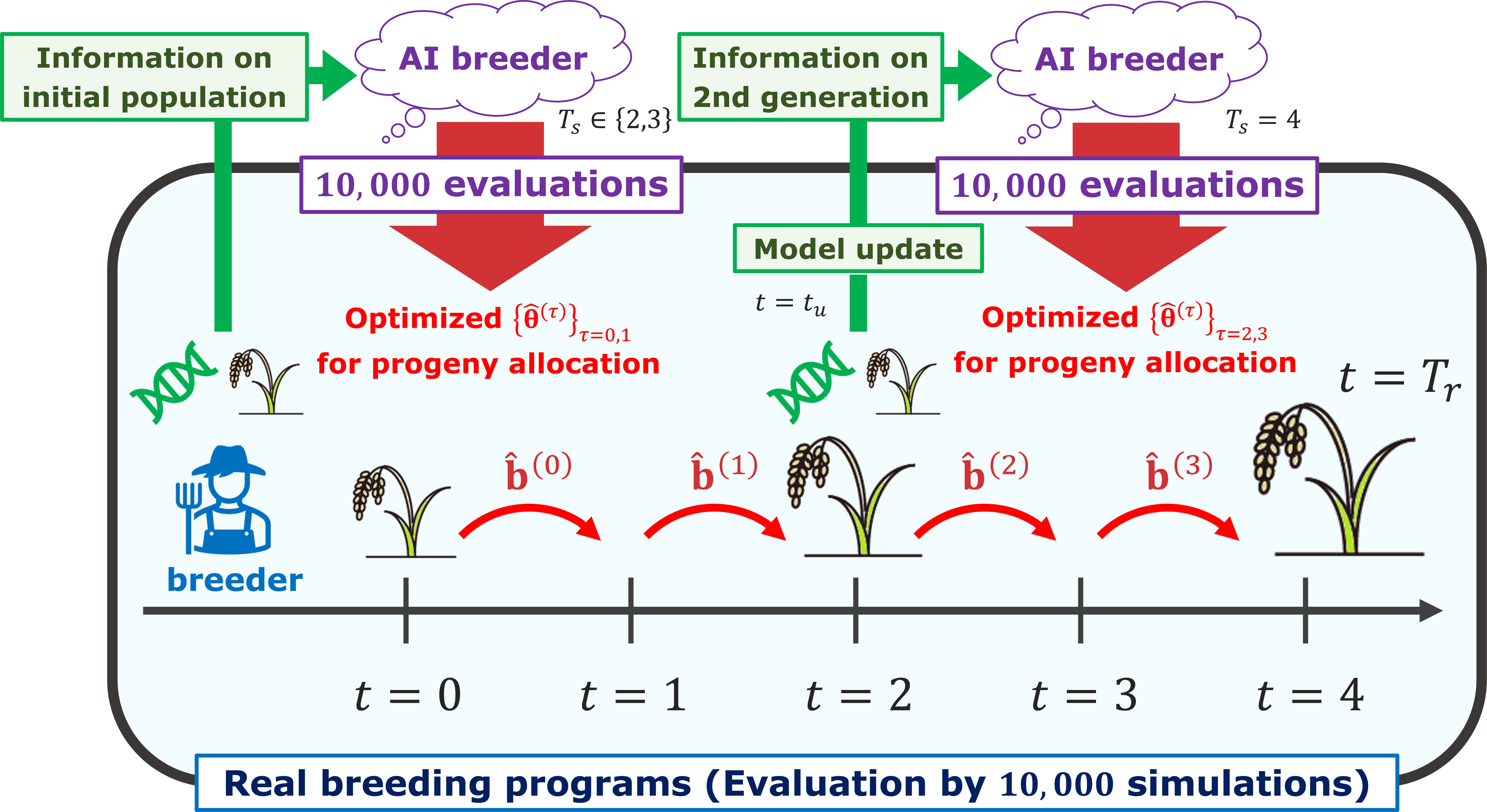
Optimization of allocation strategies in breeding schemes considering model updates and their implementation in real breeding programs. In this study, we not only estimate marker effects and optimize allocation strategies in the initial population, but also examine updating the GP model and re-optimizing allocation strategies at an intermediate stage (*t* = 2) as the breeding scheme progresses under the optimized allocation strategy. 17.4 cm (W) × 9.66 cm (H)

### Simulation of the initial breeding population and genome structure

The marker genotype and QTLs of the initial breeding population were simulated following Hamazaki and Iwata (2024). Using the coalescent simulator GENOME (Liang et al., 2007), we first generated candidate loci for genome-wide markers and QTLs of founder haplotypes. We assumed a virtual diploid crop with ten chromosomes (2*n* = 20), with 500 loci (including both markers and QTLs) on each chromosome. First, we independently generated 4,000 founder haplotypes per chromosome using GENOME with these parameters: “-pop 1 4000 -N 4000 -c 1 -pieces 100000 -rec 0.0001 -s 4000 - maf 0.01 -tree 0 -mut 0.00000001”. From these 4,000 loci, we randomly selected 500 loci with a minor allele frequency (MAF) ≥ 0.01. We then created a historical population of 2,000 founder genotypes by sampling two haplotypes with replacement from all founder haplotypes, as implemented in the R package “BreedingSchemeLanguage” by Yabe et al. (2017). This historical population then underwent random mating for 300 generations, followed by selecting 200 genotypes that were randomly mated for 15 generations under the assumption of a population bottleneck to create extensive linkage disequilibrium. Finally, we expanded the population to 2,000 genotypes and conducted random mating for three generations to produce 2,000 pre-breeding materials, following the methodology of Müller et al. (Müller et al., 2017, 2018).

Following the creation of pre-breeding materials, we selected *N* = 250 representative genotypes for the initial breeding population to capture the full genetic diversity of the pre-breeding materials, as described in Hamazaki and Iwata (2024). To ensure comprehensive representation, we employed the k-medoids clustering method using the “pam” function of the R package “cluster” version 2.1.2 (Maechler et al., 2021).

This approach divided the pre-breeding materials into 250 distinct groups based on their genotype scores, from which we selected the medoid of each group as a representative genotype. Throughout this study, genotype scores were coded as: 0 for homozygous reference alleles, 1 for heterozygous alleles, and 2 for homozygous alternative alleles. The resulting initial population had a relatively small number of genotypes, which is typical of small-scale breeding schemes where strategy optimization has a significant impact.

### Simulation of QTLs and phenotypic values

In simulating QTLs and phenotypes, we assumed one quantitative trait with a simple genetic architecture as a target trait. First, from the 500 loci simulated in the previous subsection, we randomly selected *q* loci per chromosome as QTLs. Here, we prepared two scenarios with different numbers of QTLs (i.e., *q* = 3 in Scenario 1 and *q* = 60 in Scenario 2). For simplicity, all QTL effects were additive and independently sampled from the same normal distribution. Then, we calculated true additive genotypic values (breeding values) for the initial population by multiplying QTL genotype scores by their effects and summing the products (Meuwissen *et al*., 2001). We then generated phenotypic values by adding random errors to these genotypic values. These errors were independently sampled from the identical normal distribution, whose variance was determined assuming a heritability of 0.7 throughout the study.

### Genomic prediction models and selection criteria

In this study, the AI breeder could not directly access QTL information or true genotypic values. Instead, it constructed a GP model using genome-wide markers and phenotypic data to estimate marker effects. Thus, we randomly selected 400 non-QTL loci per chromosome as markers (totaling *M* = 4,000 markers) to build this GP model. In this study, a Bayesian linear regression model in Equation 3 was used as the GP model (Meuwissen et al., 2001).

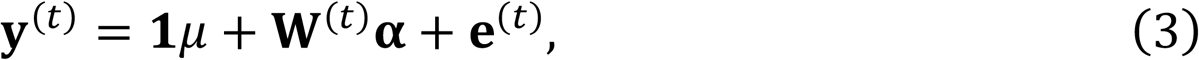

where **y**^(*t*)^ ∈ ℝ^*N*^ is a vector of phenotypic values at generation *t*, **1** ∈ ℝ^*N*^ is a vector that vertically aligns 1, *μ* ∈ ℝ is an intercept, **W**^(*t*)^ ∈ {0,1,2}^*N*×*M*^ is a matrix of marker genotype scores at generation *t*, **α** ∈ ℝ^*M*^ is a vector of marker effects, and **e**^(*t*)^ ∈ ℝ^*N*^ is an error term. Here, by appropriately designing the prior distribution of marker effects, various Bayesian linear regression models can be implemented. For this study, we used the BayesB model (Meuwissen et al., 2001; Heffner et al., 2011b) for Scenario 1 with a small number of QTLs and the Bayesian Ridge Regression model (Meuwissen et al., 2001) for Scenario 2 with a large number of QTLs. These models were implemented using the R package “BGLR” version 1.0.8 (Pérez and de los Campos, 2014).

### Breeding schemes with and without model updates

The breeding scheme assumed in this study closely followed the one described by Hamazaki and Iwata (2024), except for the updates to GP models and allocation strategies conducted during the breeding process. For simplicity and due to computational limitations, we consistently employed a small-scale breeding scheme with simple recurrent genomic selection targeting minor crops such as NUS. The key difference from previous research was that this study contained several discrepancies between the breeding schemes in the real-world context (Fig. 2A) and those simulated by the AI breeder (Fig. 2B).

1. Set initial and final generation numbers To start a breeding scheme, we set the current and final generation numbers as *t* = 0 and *T*_*r*_ = 4 for the scheme conducted by actual breeders (Fig. 2A). In contrast, for the breeding simulations by the AI breeder, the current generation was initialized as *τ* = *τ*_0_, where *τ*_0_ ∈ {0, *t*_*u*_} represented an initial generation number in breeding simulations in the scheme with model updates (Fig. 2B). In other words, the AI breeder optimized allocation strategies at *t* = 0 and *t* = *t*_*u*_, coinciding with the GP model updates (Supplemental Fig. 1). The final generation in the breeding simulations was set as *T*_*s*_ ∈ {2,3} for *τ*_0_ = 0 and *T*_*s*_ = 4 for *τ*_0_ = *t*_*u*_. This means that during the initial optimization process, we prepared two strategies depending on the target generation (*T*_*s*_ ∈ {2,3}), whereas during the second optimization of progeny allocation, we consistently targeted the final generation in the real-world breeding scheme (*T*_*s*_ = *T*_*r*_ = 4). For the comparative breeding scheme without model updates, the AI breeder optimized progeny allocation starategy only once, at the initial generation (*t* = 0) (Supplemental Fig. 2).
2. Construct GP models and optimize progeny allocation strategies At generation *t* = 0 in the scheme implemented by actual breeders, marker effects were estimated based on the marker genotype data and phenotypic data of the initial population, as described in the subsection *Genomic prediction models and selection criteria*. The progeny allocation strategies for subsequent generations were also optimized based on this initial population information, as outlined in the subsectoin *Allocation strategies to be optimized*. For breeding schemes with model updates, both GP model construction and optimization of allocation strategies were conducted again at generation *t* = *t*_*u*_, where *t*_*u*_ = 2 throughout the study (Fig. 1 and Fig. 2A). However, no model updates were implemented in the breeding simulations performed by the AI breeder (Fig. 2B).
3. Select parent candidates From the current population at generation *t*, we selected *n* genotypes as parent candidates for mating based on the increasing order of weighted breeding values (WBVs), **u**^(WBV)(*t*)^, as given in Equation 4. WBVs are variants of breeding values designed to achieve better gains in long-term programs by emphasizing rare alleles and thus maintaining genetic diversity (Jannink, 2010).

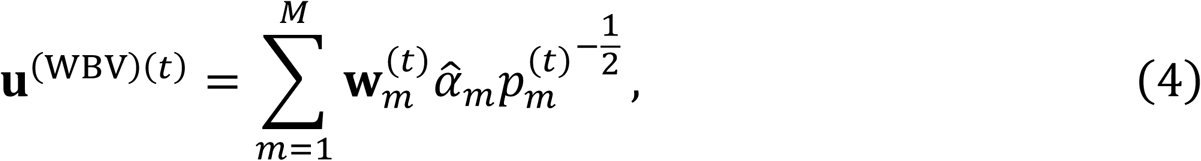 where 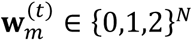 is a vector of genotype scores at marker *m* in generation *t*, 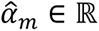 is a marker effect at marker *m*, and 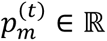 is a minor allele frequency at marker *m* in generation *t*. Here, we considered the following three methods to examine the impact of selection intensity on progeny allocation optimization.

A. SI1: Selected top *n* = 15 genotypes from the current population.
B. SI2: First performed hierarchical clustering based on marker genotypes, creating 25 clusters. Then, it selected only the top one genotype from each cluster, resulting in *n* = 25 genotypes.
C. SI3: First performed hierarchical clustering based on marker genotypes, creating 10 clusters. Then, it selected top five genotypes from each cluster, resulting in *n* = 50 genotypes.

For both SI2 and SI3, we performed hierarchical clustering using the genomic relationship matrix calculated from marker genotypes. We computed this matrix using the calcGRM function from the RAINBOWR package with version 0.1.35 (Hamazaki and Iwata, 2020) and conducted hierarchical clustering with the hclust function from the stats package with version 4.4.1 (R Core Team, 2024). This clustering procedure was identical in both the actual breeding scheme and simulations by the AI breeder.
4. Mating pair candidates for the next generation Based on the selected *n* parent candidates, mating pair candidates for the next generation *t* + 1 were determined by a half diallel crossing including selfing, resulting in 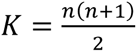 pair candidates in total.
5. Select pairs and allocate progenies to each pair Next, we employed the following three strategies for the selection of pairs and allocation of progenies. Here, to account for crossing costs, we implemented a constraint requiring that progeny allocations be made in units of at least *L* = 5 individuals per pair.

A. Optimized Resource Allocation (ORA) The first strategy was the optimized progeny allocation proposed by Hamazaki and Iwata (Hamazaki and Iwata, 2024), which directly optimized the number of progenies allocated to each pair candidate, 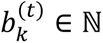, by integrating future-oriented simulations and the black-box-based optimization technique. Details are given in the *Allocation strategies to be optimized* subsection. When we assumed the breeding schemes with model updates, we considered two variants of the ORA depending on the target generation number *T*_*s*_ in the first optimization process at generation *t* = 0: ORA2 for *T*_*s*_ = 2 and ORA3 for *T*_*s*_ = 3.
B. EQual allocation (EQ) The second strategy was equal allocation, which distributed progeny evenly among mating pairs. When the number of potential mating pair candidates *K* exceeded the population size *N*, we randomly selected 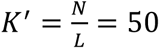 pairs from all mating pair candidates and allocated 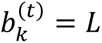 progenies to each selected pair.
C. Optimal Cross Selection (OCS) The third strategy was the OCS approach, which selected crossing pairs by considering both the GEBVs of the selected individuals and their genetic diversity (Gorjanc et al., 2018; Allier et al., 2019b). Details are provided in Supplemental Text 1, but we implemented OCS following the method in Sakurai et al. (Sakurai et al., 2024), which resulted in the selection of *K*^′′^ = 25 pairs from all mating pair candidates. After selecting these *K*^′′^ pairs, 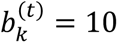 progenies were allocated to each selected mating pair.
6. Generate progeny for the next generation From mating pair *k*, *b*_*k*_ progenies were produced following the principle of meiosis, resulting in a new population with generation *t* + 1 consisting of *N* progenies. Here, recombination rates used in the meiosis simulation were calculated using the Kosambi map function (Kosambi, 1943; Zhao and Speed, 1996) applied to the linkage map generated by GENOME.
7. Repeat the selection and mating processes Steps 2-6 were repeated until reaching the population with the final generation *T*_*r*_ = 4 (or *T*_*s*_ for breeding simulations by the AI breeder), assuming a scenario where rapid genetic improvement was needed in small-scale breeding schemes.

**Fig. 2.**
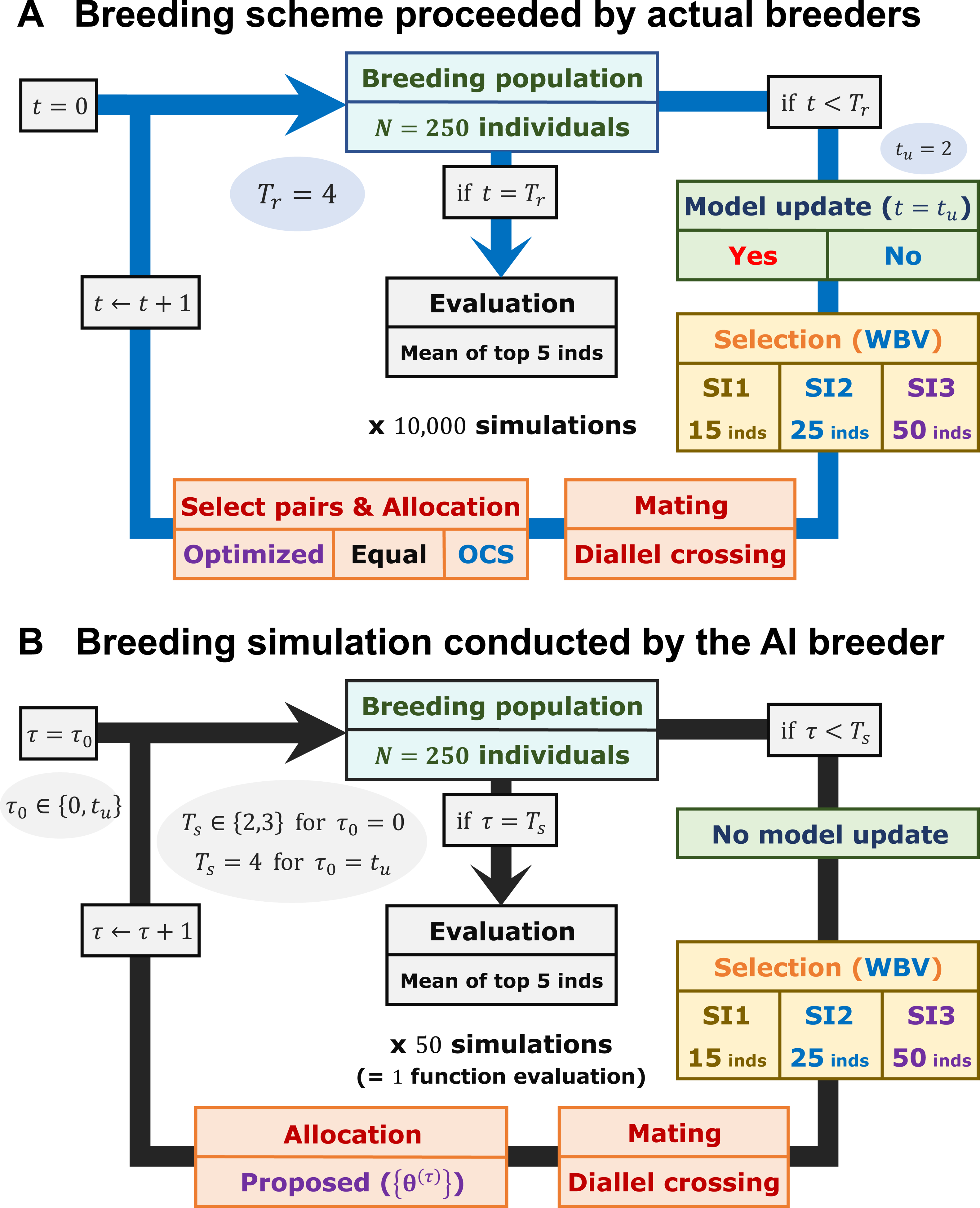
Images of breeding schemes assumed in this study. Similar to Hamazaki and Iwata (Hamazaki and Iwata, 2024), we assumed small-scale plant breeding programs with recurrent selection. In a scheme, the selection, mating, and allocation steps were repeated until the final generation. Here, we prepared for the two scenarios with different numbers of QTLs and three selection strategies with different selection intensities. (A) Breeding schemes proceeded by actual breeders for the evaluation of each strategy. (B) Breeding simulations conducted by the AI breeder used for the optimization of allocation strategies in ORA variants. 17.4 cm (W) × 20.63 cm (H)

### Evaluation of the genetic gains of breeding schemes

We evaluated the results for each strategy using breeding values at generation *t*, i.e., **u**^(*t*)^ = **W**^(*t*)^**α**, particularly for the final population with generation *T*_*r*_. Here, to evaluate the population’s maximum under the breeding strategy, we computed 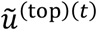, the empirical mean of **u**^(*t*)^ for the top five genotypes. These genotypes were selected based on their **u**^(*t*)^ values in descending order. We then scaled 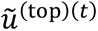 by the initial population’s mean and standard deviation, *u*^(mean)(0)^ and 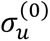, resulting in 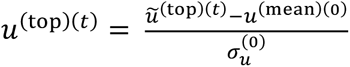. We selected the top five genotypes for evaluation to represent potential for new variety development while mitigating stochastic variation between simulations. For each parent panel simulation dataset, we ran 10,000 breeding schemes per strategy and calculated the genetic gain for evaluation as the empirical mean of *g*^(*t*)^ = *u*^(top)(*t*)^/*u*^(top)(0)^ from these results. We repeated the simulation and evaluation processes for 10 replicates of the phenotype simulation, each with different QTL positions and effects.

### Optimization of progeny allocation strategies

In this study, the number of progenies allocated to pair *k* at generation *τ*, 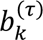, was determined by applying the softmax function to the weighted sum of the multiple “goodness” metrics (Supplemental Fig. 3), following Hamazaki and Iwata (Hamazaki and Iwata, 2024). Since we need to incorporate the constraint on the minimum allocation units *L*, 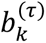 was computed as shown in Equation 5.

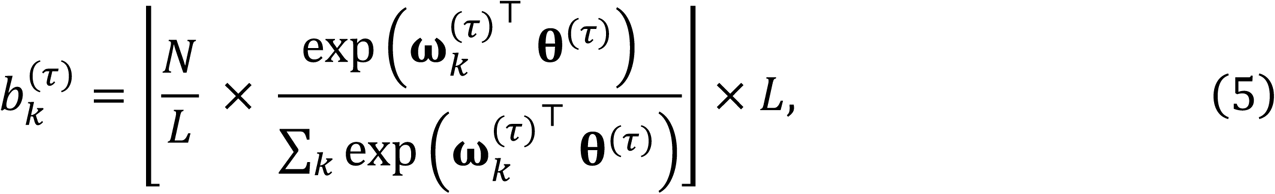

where 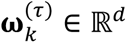 is a feature vector of goodness metrics, **θ**^(*τ*)^ ∈ ℝ^*d*^ is a vector that determines the weight given to each goodness metric, thus becoming a parameter to be optimized by the AI breeder, and ⌊*x*⌋ is the floor function of *x*. Here, as in Equations 1 and 2, the breeding scheme can be viewed as a black-box function *F* with 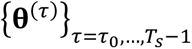 as input and the final genetic gain, 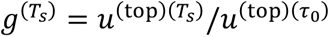, as output. Thus, we employed a black-box optimization technique called StoSOO (Valko *et al*., 2013) to estimate the weighting parameters that maximized 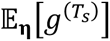, following Hamazaki and Iwata (2024). In this study, we conducted 50 breeding simulations to calculate the expectation with respect to **η** for one function evaluation and performed 10,000 or 20,000 function evaluations to obtain the optimal set of weighting parameters for breeding schemes with or without model updates, respectively (Fig. 3 and Supplemental Fig. 2). Then, the optimized 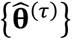 was utilized for progeny allocation in the real breeding schemes conducted by actual breeders.

**Fig. 3.**
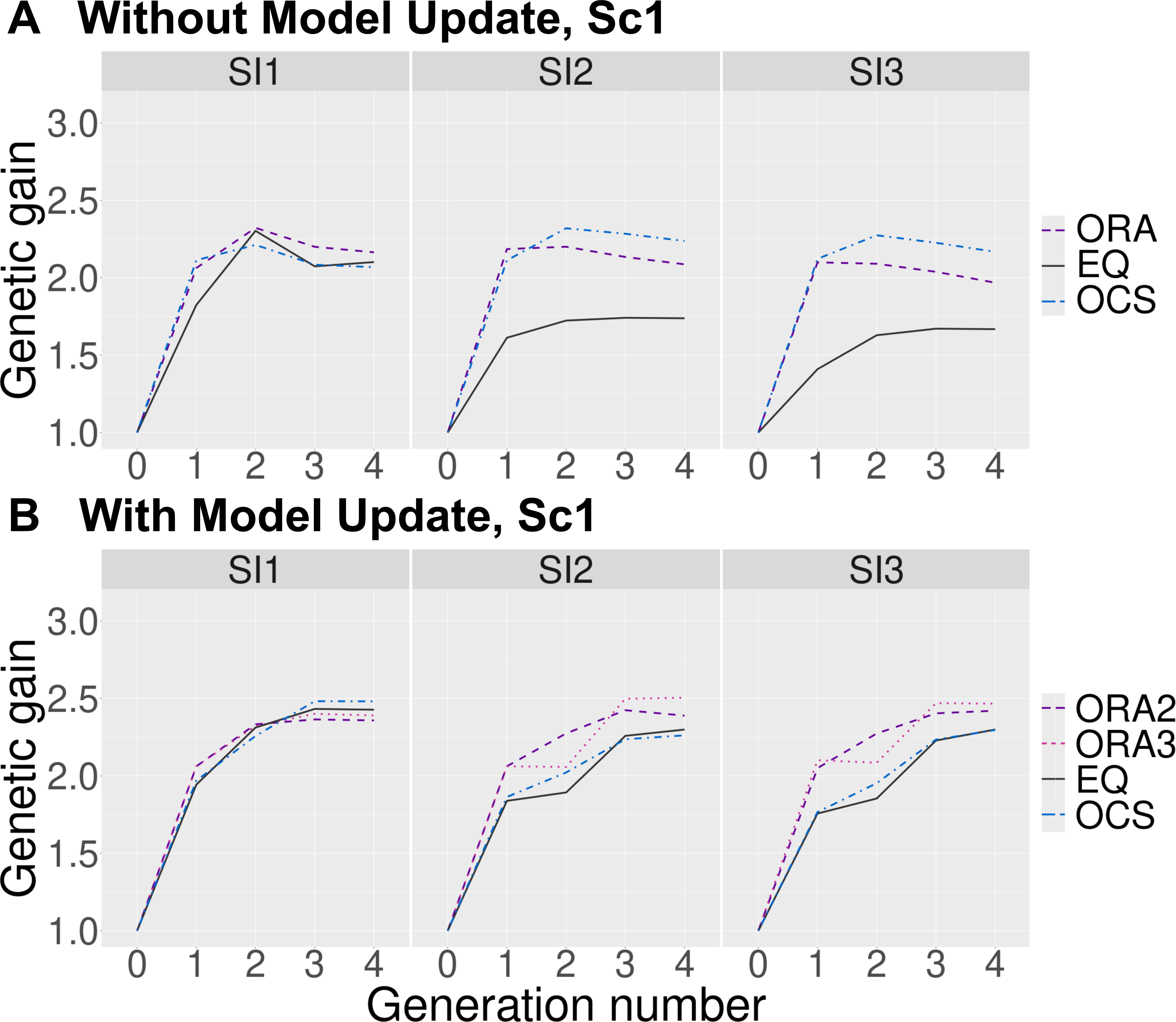
Change in the expected genetic gains over four generations under different selection intensities for Scenario 1. The horizontal and vertical axes represent the number of generations and the genetic gains, respectively. We compared the following allocation strategies under the different selection intensities (SI1 – SI3): ORA: optimized resource allocation (purple dashed), ORA2: optimized resource allocation with targeting *Ts* = 2 in the initial optimization (purple dashed), ORA3: optimized resource allocation with targeting *Ts* = 3 in the initial optimization (light purple dotted), EQ: equal allocation (black solid). OCS: optimal cross selection (blue dash-dotted line). Panel (A) corresponds to the scheme without model updates (comparing ORA, EQ, and OCS), while panel (B) corresponds to the scheme with model updates (comparing ORA2, ORA3, EQ, and OCS). 17.4 cm (W) × 14.9 cm (H)

As candidates for the feature vector of goodness metrics 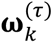, we used the WBV (Goddard, 2009; Jannink, 2010) and the expected genetic variance of progeny (GVP) for each mating pair (*d* = 2). The first metric, WBV, for pair *k* was calculated as the mean of the two parents’ WBVs in Equation 4, i.e., 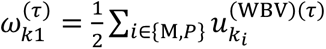, where *k*_M_ and *k*_P_ correspond to the indices of the parents. The second metric, GVP, was assessed as the genetic diversity of gametes produced after self-fertilization of progeny from each mating pair. The GVP for mating pair *k* can be theoretically derived using approaches proposed by previous studies (Lehermeier et al., 2017; Allier et al., 2019a; b), as shown in the following Equation 6.

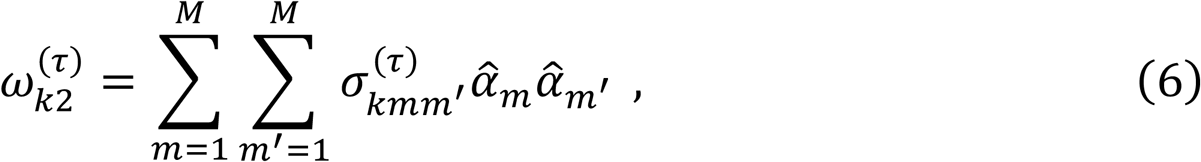

where 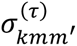 represents the covariance between markers *m* and *m*^′^ resulting from progeny segregation derived from pair *k* at generation *τ*. The detailed derivation of 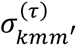 is provided in Allier et al. (2019b), but 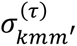 can be computed as a function of allelic scores of the parents’ haplotypes and the recombination rates between markers. Here, when markers *m* and *m*^′^ are located on different chromosomes, 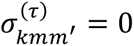, and this term does not contribute to the GVP.

## Results

### Change in the genetic gains over four generations

We compared three allocation strategies (ORA, EQ, and OCS) in breeding schemes without model updates, and four strategies (ORA2, ORA3, EQ, and OCS) in schemes with model updates. First, we plotted the genetic gains over four generations across two scenarios for the number of QTLs (Scenario 1 and Scenario 2) under three selection intensities (SI1, SI2, and SI3) (Fig. 3 and Supplemental Fig. 4).

In the breeding schemes without model updates, genetic gains reached saturation at generation *t* = 2 for all allocation strategies across both scenarios and all selection intensities (Fig. 3A and Supplemental Fig. 4A). In Scenario 1 with a small number of QTLs, comparative analysis of allocation strategies revealed that ORA exhibited superior performance over both OCS and EQ under high selection intensity (SI1), whereas under lower selection intensities (SI2, SI3), the genetic gains were maximized by OCS, followed by ORA and EQ (Fig. 3A). Conversely, in Scenario 2 with a large number of QTLs, ORA demonstrated genetic gains that were either equal to or greater than OCS across all selection intensities, while consistently outperforming EQ (Supplemental Fig. 4A). Given that genetic gains quickly plateaued across all allocation strategies, implementing model updates are necessary for developing more accurate breeding strategies.

When examining breeding schemes with model updates, genetic gains continued to improve after the model update at generation *t* = 2, progressing until the final generation (*t* = 4) across all scenarios, selection intensities, and allocation strategies (Fig. 3B and Supplemental Fig. 4B). This trend was particularly pronounced in Scenario 2, suggesting that model updates are highly effective for quantitative traits governed by a large number of QTLs (Supplemental Fig. 4B). In Scenario 1 with fewer QTLs (Fig. 3B), ORA strategies performed equally or worse compared to EQ and OCS under high selection intensity (SI1), yet exhibited superior genetic gains under lower selection intensities (SI2, SI3). A similar trend was observed in Scenario 2 with numerous QTLs (Supplemental Fig. 4B); however, there were two notable differences from Scenario 1: ORA3 demonstrated superior performance compared to other allocation strategies even under SI1, and OCS clearly outperformed EQ under SI2 and SI3, although it still could not match ORA’s performance. When comparing the two ORA strategies, ORA3 consistently demonstrated superior performance over ORA2, although the difference was small, except in Scenario 2 under SI3 (Fig. 3B and Supplemental Fig. 4B).

### Genetic gains across different simulation repetitions

To evaluate the characteristics of each allocation strategy comprehensively, we analyzed the cumulative distribution functions (CDFs) of genetic gain in the final generation (*t* = 4) across two scenarios under three selection intensities, using 10,000 simulation repetitions. Figure 4 and Figure S5 show these results, with the horizontal axis indicating final genetic gain and the vertical axis indicating the percentile of simulation repetitions. The 1st and 99th percentiles represent the worst and best performances among the 10,000 repetitions. Effectiveness of each strategy is indicated by how far its CDF curve shifts rightward and downward—the more significant this shift, the better the overall performance of the strategy.

**Fig. 4.**
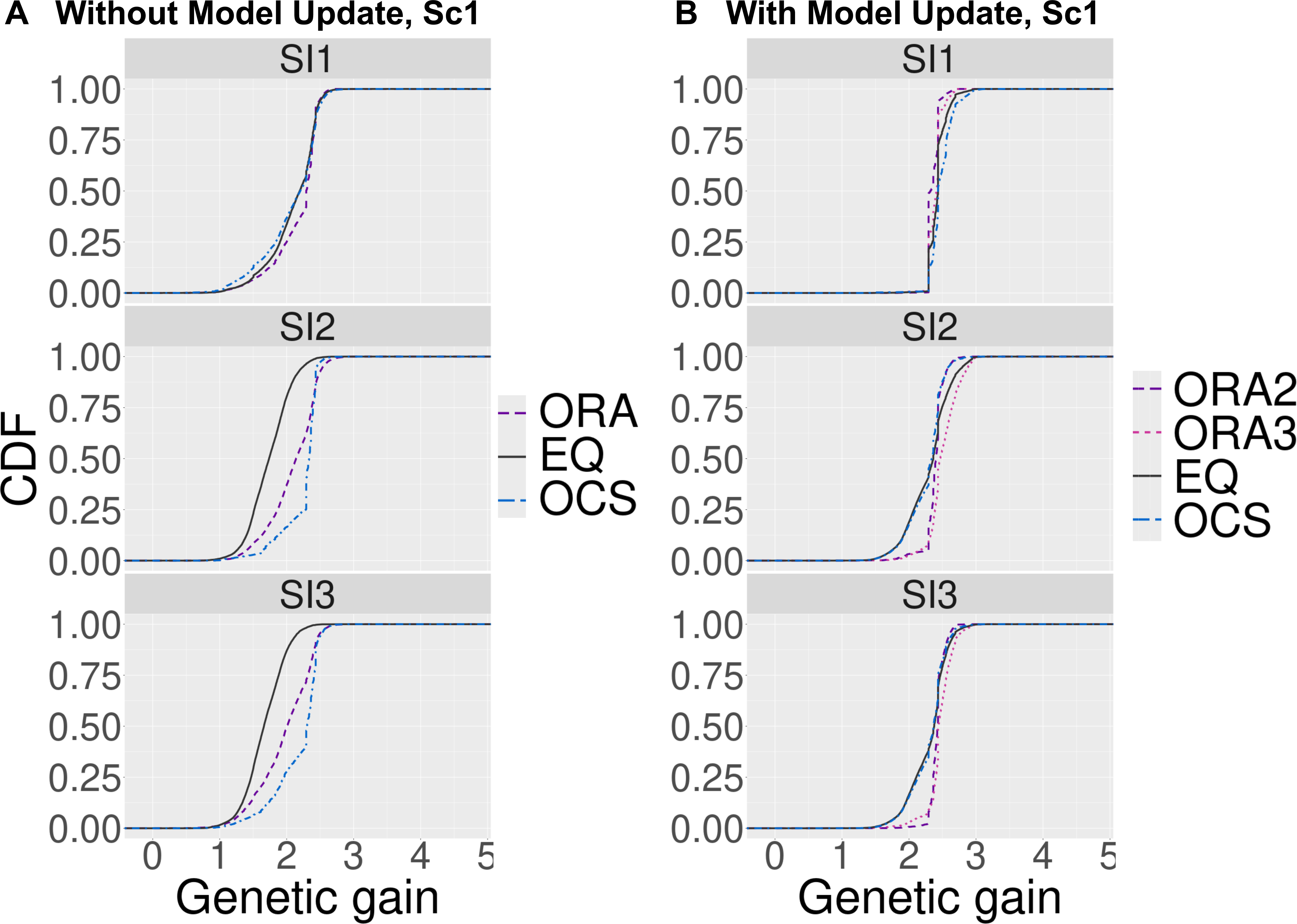
Genetic gains across different simulation repetitions for Scenario 1. The horizontal axis shows the final genetic gain of an individual, whereas the vertical axis represents the percentile of the simulation repetitions. Panel (A) corresponds to the scheme without model updates (comparing ORA, EQ, and OCS), while panel (B) corresponds to the scheme with model updates (comparing ORA2, ORA3, EQ, and OCS). The abbreviations of the allocation strategies are the same as those of Fig. 3. 17.4 cm (W) × 12.49 cm (H)

Regarding the comparison between allocation strategies, the CDF curves showed similar trends to those observed in Fig. 3 and Supplemental Fig. 4 (Fig. 4 and Supplemental Fig. 5). For both schemes with and without model updates, the CDF curves for the ORA strategies in Scenario 2 (Supplemental Fig. 5) showed smoother and gentler slopes than those in Scenario 1 (Fig. 4), suggesting that the optimization for traits governed by numerous QTLs led to greater variations among simulation repetitions. In Scenario 1 with model updates, the CDF curves exhibited slopes almost parallel to the vertical axis for all allocation strategies under high selection intensity, and for the optimized allocation strategies under low selection intensities (Fig. 4B). These results indicated that applying strong selection intensity to traits controlled by a small number of QTLs leads to consistent outcomes across different simulation runs. In addition, in Scenario 2 with model updates, while the ORA strategies performed comparably to OCS in the best-performing case, it consistently outperformed OCS in the worst-performing case, indicating that they can provide more stable results for traits controlled by numerous QTLs (Supplemental Fig. 5B).

### Genetic gains across different phenotype simulations

We further evaluated genetic gains in the final generation (*t* = 4) using ten replications of phenotype simulations to verify that the ORA strategies outperformed the EQ and OCS strategies across traits with varying QTL positions, effects, and errors. For these simulations, we changed QTL positions and effects while maintaining consistent QTL numbers and effect distributions. To reduce computational time, we determined the ORA strategies after 2,500 function evaluations for schemes with model updates and 5,000 for schemes without. The expected genetic gains for all ten replications appear in Supplemental Fig. 6 and Supplemental Fig. 7, while the improvement rates compared to EQ, calculated by subtracting the genetic gains of EQ from those of ORA or OCS and then dividing by the genetic gains of EQ, are presented in Fig. 5 and Supplemental Fig. 8.

**Fig. 5.**
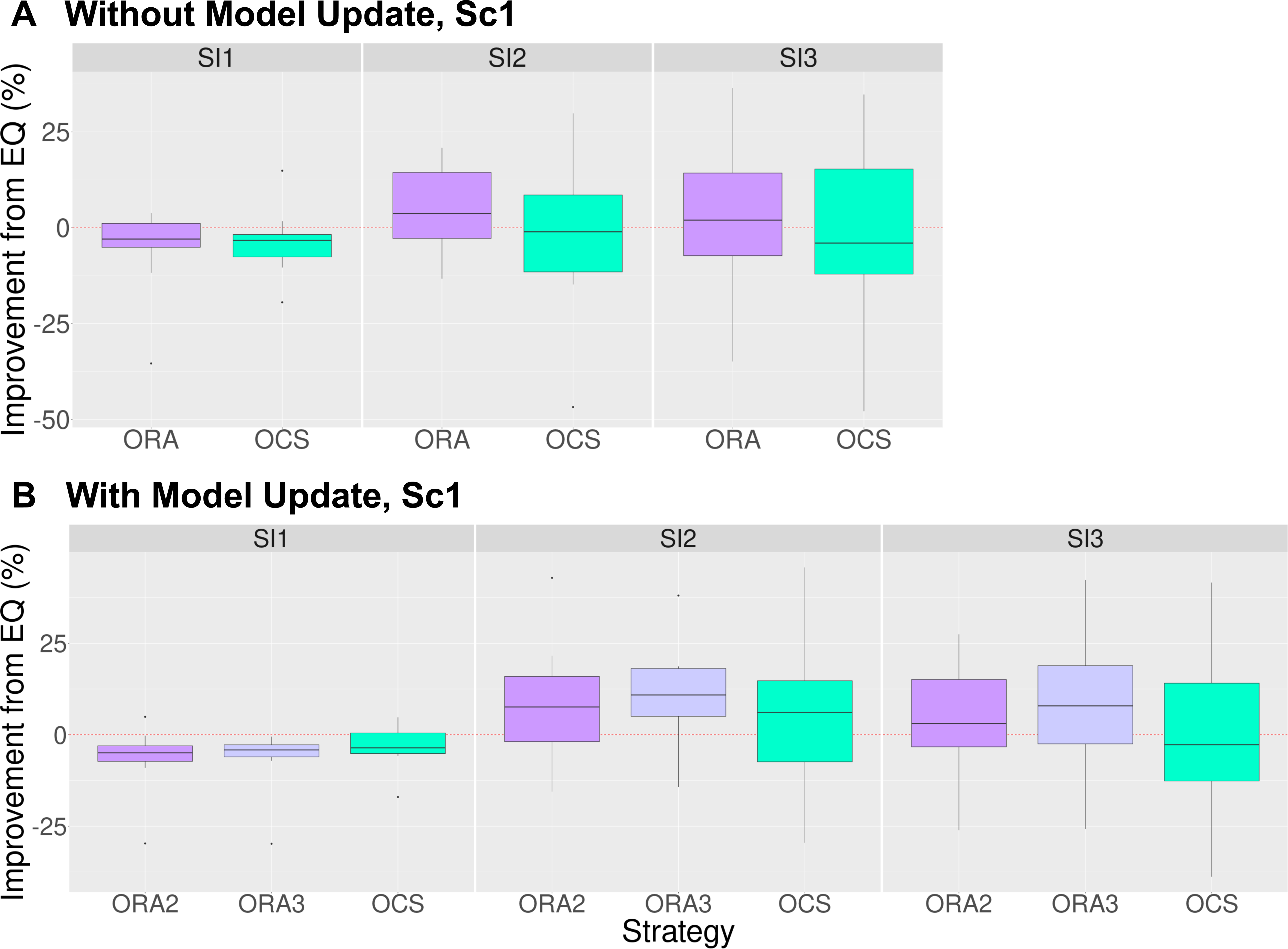
Improvement of the final genetic gains compared to the equal allocation strategy using ten replications for the phenotype simulation under different selection intensities for Scenario 1. This figure includes boxplots of the improvement of the final genetic gains compared to the equal allocation strategy using ten replications for the phenotype simulation. The horizontal and vertical axes represent the different allocation strategies and the improvement rate of the final genetic gains, respectively. We compared the following allocation strategies under the different selection intensities (SI1 – SI3): ORA (purple), ORA2 (purple), ORA3 (light purple), and OCS (light blue), with details given in Fig. 3. Panel (A) corresponds to the scheme without model updates (comparing ORA and OCS), while panel (B) corresponds to the scheme with model updates (comparing ORA2, ORA3, and OCS). 17.4 cm (W) × 12.98 cm (H)

While the ORA strategies were unable to outperform EQ under high selection intensity, they were markedly superior to EQ under low selection intensities (Fig. 5 and Supplemental Figs 6 – 8). This trend was particularly pronounced when model updates were implemented in Scenario 2 with a large number of QTLs (Supplemental Fig. 7B and Supplemental Fig. 8B). In addition, comparing the ORA strategies with OCS revealed that in Scenario 2, ORAs failed to match or exceed OCS, regardless of whether model updates were implemented (Supplemental Fig. 8). However, in Scenario 1, ORA strategies surpassed OCS in improvement rates (Fig. 5). These findings suggest that the ORA strategies are particularly effective for traits controlled by a small number of QTLs. Furthermore, in Scenario 2, where traits are governed by numerous QTLs, OCS exhibited greater variability across different phenotype simulations, indicating that ORA might be more effective in developing varieties with consistent genetic gains across diverse traits (Supplemental Fig. 8).

### Change in the genetic variances over four generations

We then compared genetic diversity between ORA, EQ, and OCS strategies across four generations, examining schemes both with and without model updates under three selection intensities for Scenario 1 (Supplemental Fig. 9) and Scenario 2 (Supplemental Fig. 10). Genetic diversity within each generation’s breeding population was measured using the genetic variance of true genotypic values, i.e., 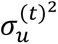 described in the subsection *Evaluation of the genetic gains of breeding schemes*.

In Scenario 1, genetic variance increased compared to the initial population until generation *t* = 2, regardless of whether model updates were implemented (Supplemental Fig. 9). Subsequently, breeding populations without model updates maintained stable genetic variance (Supplemental Fig. 9A). In contrast, when model updates were performed, genetic variance declined for all strategies under SI1, and for all strategies except EQ under SI2 and SI3 (Supplemental Fig. 9B). In Scenario 2, both ORA and EQ strategies showed slight increase in genetic variance regardless of model updates, while OCS exhibited a gradual decrease (Supplemental Fig. 10). When comparing allocation strategies in the final generation, ORA maintained lower genetic variance than EQ (except in Scenario 1 without model updates), but preserved higher genetic variance than OCS.

### Change in the prediction accuracies over four generations

Finally, we assessed prediction accuracy across four generations by calculating the correlation between true and estimated genotypic values in each generation for Scenario 1 (Fig. 6) and Scenario 2 (Supplemental Fig. 11).

**Fig. 6.**
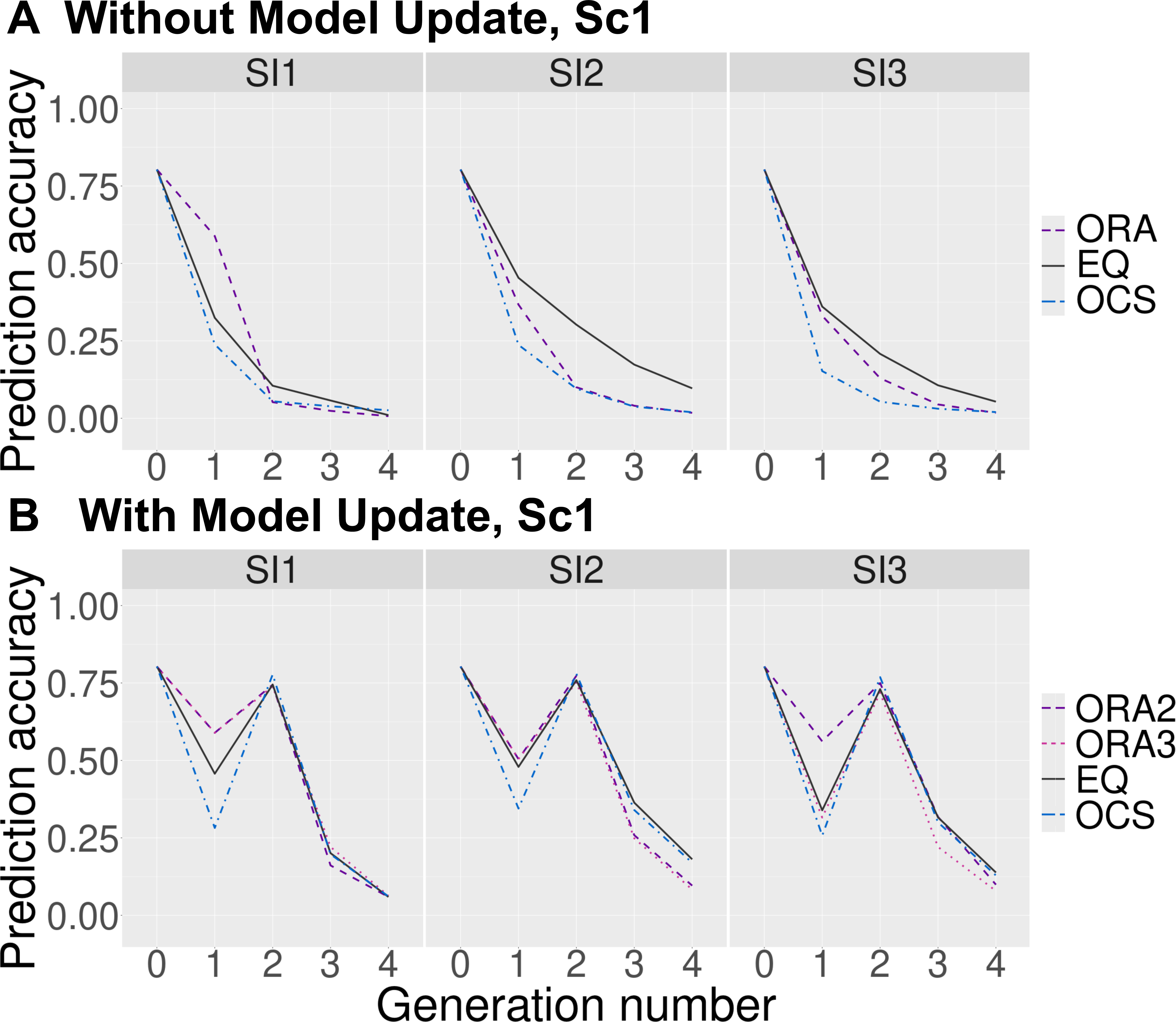
Change in the prediction accuracies over four generations under different selection intensities for Scenario 1. The horizontal and vertical axes represent the number of generations and the prediction accuracies for the breeding populations, respectively. Panel (A) corresponds to the scheme without model updates (comparing ORA, EQ, and OCS), while panel (B) corresponds to the scheme with model updates (comparing ORA2, ORA3, EQ, and OCS). The abbreviations of the allocation strategies are the same as those of Fig. 3. 17.4 cm (W) × 14.9 cm (H)

First, without model updates, prediction accuracy declined to nearly zero by the final generation (*t* = 4) in both scenarios across all selection intensities and allocation strategies (Fig. 6A and Supplemental Fig. 11A). Most notably, except for EQ under SI2, accuracy had already fallen to approximately 0.1 by *t* = 2, indicating rapid population changes from selection and mating.

In contrast, implementing model updates at *t* = 2 restored prediction accuracy to approximately 0.75 across all scenarios, selection intensities, and allocation strategies—comparable to those in the initial population (Fig. 6B and Supplemental Fig. 11B). Although accuracy gradually decreased toward the final generation, breeding schemes with model updates maintained substantially better accuracy than those without updates.

Between scenarios, Scenario 2, with more QTLs controlling traits, showed faster decreases in prediction accuracy than Scenario 1 (Supplemental Fig. 11 compared with Fig. 6), though model updates effectively restored accuracy to similar levels in both scenarios. Among allocation strategies, OCS consistently showed the lowest prediction accuracy (Fig. 6 and Supplemental Fig. 11), particularly in Scenario 2 where accuracy plummeted to approximately 0.05 by generation *t* = 1. For ORA and EQ strategies, EQ showed faster decline in accuracy under high selection intensity (SI1), while ORA variants declined faster under moderate and low selection intensities (SI2 and SI3). Interestingly, in Scenario 2, EQ showed marked accuracy decreases under high selection intensity, whereas ORA exhivited similar pattern of accuracy decline across all selection intensities (Supplemental Fig. 11). Between ORA variants, ORA3 showed slightly greater reduction in accuracy than ORA2, though differences were minimal.

## Discussion

Breeding schemes involve multiple decision-making points, such as selection and mating. Recently, numerous breeding strategies leveraging genomic information have emerged (Wang et al., 2018; Moeinizade et al., 2019; Diot and Iwata, 2022; Hamazaki and Iwata, 2024; Sakurai et al., 2024). Previous research, including work by Hamazaki and Iwata (2024), often developed and optimized breeding strategies assuming that true QTL effects are known (Kemper et al., 2012; Daetwyler et al., 2015; Goiffon et al., 2017; Wang et al., 2018; Moeinizade et al., 2019, 2022; Amini et al., 2021; Zhang and Wang, 2022; Hamazaki and Iwata, 2024; Hamazaki et al., 2024). However, in practical breeding schemes, true QTL effects are seldom fully understood, requiring the use of marker effects estimated by GP instead. Thus, in this study, we optimized allocation strategies using estimated marker effects while incorporating model updates, and subsequently evaluated their effectiveness. The following discussion examines our findings on genetic gains and genetic variances, and assesses whether optimized allocation strategies remained effective when using estimated marker effects rather than true QTL effects.

### Discussion on genetic gains

First, examining the changes in genetic gains revealed that while genetic gain plateaued without model updates (Fig. 3A and Supplemental Fig. 4A), implementing model updates consistently improved gains through the final generation (Fig. 3B and Supplemental Fig. 4B). This improvement is strongly linked to enhanced prediction accuracy from these updates (Fig. 6B and Supplemental Fig. 11B), as making selection, mating, and allocation decisions using more accurate models allowed genetic gain to increase through the final generation. Notably, the improvement between generations *t* = 2 and *t* = 3 exceeded that between *t* = 1 and *t* = 2, likely because the prediction accuracy recovered after model updates at *t* = 2, enabling more effective selection.

Our analysis revealed interesting patterns in the relationship between selection intensity and allocation strategy (Figs. 3 – 5 and Figs. S4 – S8). When selection intensity was high (SI1), the optimized allocation strategy (ORA) achieved genetic gains similar to or lower than those of equal allocation (EQ). However, under lower selection intensity conditions (SI2, SI3), ORA significantly outperformed EQ. This finding supports the conclusion by Hamazaki and Iwata (2024) that optimizing allocation strategy becomes more crucial when selection intensity is low.

Additionally, under low selection intensity conditions, model updates enabled ORA to achieve higher genetic gain than both EQ and OCS in both scenarios (Figs. 3 and 4 and Figs. S4 and S5). This demonstrates that ORA functions effectively with model updates because allocation strategies are re-optimized simultaneously with the models. When comparing ORA2 and ORA3, ORA3 generally produced superior results, suggesting that optimization strategies focused on generations beyond the next model update are more effective than those targeting only the immediate next model update (Figs. 3B – 5B and Figs. S4B – S8B). Consistent with this observation, ORA3, with its long-term approach, demonstrated a modest advantage in maintaining genetic variance compared to ORA2 (Figs. S9B and S10B).

### Discussion on genetic variances

In this study, genetic variances tented to increase in the first two generations for all allocation strategies (Figs. S9 and S10). While selection would generally be expected to reduce genetic diversity, this counterintuitive result stems from the experimental conditions in this study. In this study, the initial breeding population was assumed to be close to germplasm collections, so rare alleles might also be assigned genetic effects. Therefore, in the initial stages, these rare alleles were favored, shifting allele frequencies toward 0.5 and thereby contributing to the increase in genetic variances—a phenomenon observed in other studies as well (Jannink, 2010). It should be noted that in simulations using populations derived from bi-parental or four-way crosses as initial populations, as in Sakurai et al. (2024), allele frequencies generally start close to 0.5 rather than 0, preventing such increases from occurring.

With model updates implemented, subsequent generations showed different patterns in genetic variances. In Scenario 1 with fewer QTLs, genetic variances tended to decrease as alleles became fixed and minor allele frequencies approached 0 (Supplemental Fig. 9B). In contrast, Scenario 2 with more QTLs showed smaller decreases in genetic variances because many alleles remained that were not moving toward fixation (Supplemental Fig. 10B). Although our simulations spanning four generations did not reveal large decreases in genetic variance, we expect genetic variances to decrease rapidly in both scenarios during longer-term breeding schemes. In simulations without model updates, prediction accuracies in the later stages of breeding schemes were inherently low, causing individuals to be selected more randomly, which tended to suppress the decrease in genetic variances (Figs. S9A and S10A).

### Comparison of ORA and OCS

From the results of different simulation repetitions, while ORA and OCS showed similar performance in the best-performing cases, ORA significantly outperformed OCS in the worst cases (Fig. 4 and Supplemental Fig. 5). This indicates that ORA can consistently deliver high-quality varieties even when random factors such as recombination act unfavorably.

Moreover, from the results of different phenotype simulations, particularly in Scenario 1 assuming traits controlled by a small number of QTLs, ORA variants showed better genetic gains than OCS at lower selection intensities (Fig. 5 and Supplemental Fig. 6). In contrast, for Scenario 2 assuming traits controlled by many QTLs, ORA showed no clear advantage over OCS (Figs. S7 and S8). However, ORA showed less variation in results, suggesting it may have stability that can accommodate various traits. Additionally, since this study terminated the black-box optimization for ORA prematurely due to limitations in computational time, it is possible that spending more time on optimization could surpass OCS. Indeed, focusing on the results for one phenotype simulation confirmed the advantage of ORA in both scenarios (Figs. 3 and 4 and Figs. S4 and S5).

Furthermore, while OCS achieved high genetic gains through strong selection pressure, evidenced by the sharp decrease in prediction accuracy from *t* = 0 to *t* = 1, ORA achieved comparable genetic gains while better controlling rapid changes in population structure than OCS (Fig. 6 and Supplemental Fig. 11). As confirmed when updating models in Scenario 2, ORA variants effectively balanced the overall selection intensity across both selection and allocation stages through its optimized allocation strategy, shown by consistent prediction accuracy changes regardless of selection intensity (Supplemental Fig. 11B). Although this study focused on short-term breeding schemes and did not observe significant decreases in genetic variances, in longer-term breeding schemes, this appropriate adjustment of selection intensity is expected to contribute to improving genetic gain while maintaining genetic diversity.

Based on these results, ORA proves to be a robust approach that not only demonstrates excellent performance but also maintains stability when faced with diverse target traits or random factors such as recombination.

### Future prospects and conclusions

Our study demonstrates that the optimized allocation strategies, when combined with appropriate model updates, deliver excellent performance and robustness in realistic breeding schemes that rely on estimated marker effects. By conducting simulations under more realistic assumptions, our study bridges the gap between theoretical approaches and practical breeding applications, making the optimization framework valuable for plant breeders in their daily decision-making. We believe this methodology can be readily implemented by breeding programs worldwide to enhance genetic gain while maintaining diversity.

However, we found that these strategies become less effective under high selection intensity. In future works, we plan to address this limitation by optimizing selection intensity and allocation strategy simultaneously. Additionally, since model updates have enabled longer-term breeding programs, future challenges include performance verification in long-term breeding schemes and application of the optimized allocation strategies in real-world breeding schemes with field trials.

Furthermore, we aim to develop more efficient methods for optimizing breeding decisions by leveraging advanced technologies like automatic differentiation as proposed in Hamazaki et al. (2024). Through these efforts, the integration of breeding simulations and optimization approaches as proposed by Hamazaki and Iwata (2024), extended in our study via updates of the GP models and the optimal allocation strategies, will bring significant innovation in genome-assisted breeding.

## Author Contribution Statement

KH, HI, and KT designed the study; KH implemented all the resource allocation strategies; KH performed the simulation study; KH interpreted results and drafted the manuscript; KT supervised the study. All authors read and approved the final manuscript.

## Acknowledgments

This work was supported by JST, ACT-X Grant Number JPMJAX23CL, Japan. We would like to thank Dr. Kengo Sakurai for his assistance in implementing the OCS method.

