## Supplementary Data 1 for "Optimizing progeny allocation strategies in breeding schemes while updating genomic prediction models"

**Manuscript type:** Research Paper

**Manuscript information:** 6 figures and 0 tables (11 supplemental figures)

**List of author's last names:** Hamazaki, Tsuda, and Iwata

**Running title:** Optimize progeny allocation while updating GP

**Member or nonmember of the Japanese Society of Breeding:** Kosuke Hamazaki (member, 6112)

**Supplemental Text 1**

*Details of Optimal Cross Selection*

OCS selects superior crossing pairs from all the possible combinations in the current population, considering both the GEBVs and genetic diversity of the selected individuals (Gorjanc et al., 2018; Allier et al., 2019). In this study, we implemented OCS following the method in Sakurai et al., (2024). This method solves the following optimization problem to determine the selected pairs.

$$\mathrm{argmax}_{a_k} \sum_{k=1}^{K} a_k \bar{u}_k^{(\mathrm{BV})(t)} , \quad \text{(S1)}$$

$$\text{s.t. } \gamma^{(t)} > h^{(t)}, \quad \text{(S2)}$$

where $K = \frac{n(n+1)}{2}$ is the number of total mating pairs, $a_k \in \{0,1\}$ is a dummy variable that indicates whether pair $k$ is selected ($a_k = 1$) of not ($a_k = 0$), and $\bar{u}_k^{(\mathrm{BV})(t)} \in \mathbb{R}$ is the mean of the two parents' GEBVs ( $\bar{u}_k^{(\mathrm{BV})(t)} = \frac{1}{2} \sum_{i \in \{\mathrm{M,P}\}} \mathbf{w}_{k_i}^{(t)\top} \hat{\boldsymbol{\alpha}}$ ). $\gamma^{(t)} \in \mathbb{R}$ represents the genetic diversity of all selected mating pairs, while $h^{(t)} \in \mathbb{R}$ is the genetic diversity constraint at generation $t$. First, $\gamma^{(t)}$ is defined as follows in Equation S3.

$$\gamma^{(t)} = 1 - \mathbf{c}^{\top}\mathbf{K}^{(t)}\mathbf{c}. \quad \text{(S3)}$$

Here, $\mathbf{c} \in \{0,1\}^n$ is a contribution vetor to each individual, computed as $\mathbf{c} = \frac{1}{2K''}\mathbf{Za}$, where $\mathbf{Z} \in \{0,1\}^{n \times K}$ is a design matrix indicating the correspondence between parent candidates and pairs, $\mathbf{a} = [a_1, \ldots, a_K]^{\top}$, and $K'' = 25$ is the number of pairs selected by

OCS. $\mathbf{K}^{(t)} \in \mathbb{R}^{n \times n}$ is an identical-by-state matrix, which can be computed as shown in Equation S4.

$$\mathbf{K}^{(t)} = \frac{1}{2}\left(\frac{1}{M}\left(\mathbf{W}^{(t)} - 1\right)\left(\mathbf{W}^{(t)} - 1\right)^{\top} + 1\right) \tag{S4}$$

Next, $h^{(t)}$ is defined as shown in Equation S5, which was originally proposed by Allier et al. (2019).

$$h^{(t)} = \begin{cases} h^{(0)} + \left(\frac{t}{t^{\star}}\right)^{s}\left(h^{\star} - h^{(0)}\right) \text{ for } t \leq t^{\star} \\ h^{\star} \text{ for } t > t^{\star} \end{cases}, \tag{S5}$$

Here, $h^{(0)}$ is the mean heterozygosity rate of the initial population, and $t^{\star} \in \mathbb{N}$, $h^{\star} \in \mathbb{R}$, and $s \in \mathbb{R}$ are hyperparameters that define the OCS. $h^{(0)}$ is computed as shown in the following Equation S6.

$$h^{(0)} = \frac{1}{M}\sum_{m=1}^{M} 2p_m^{(0)}\left(1 - p_m^{(0)}\right), \tag{S6}$$

where $p_m^{(0)} \in \mathbb{R}$ is the minor allele frequency at marker $m$ in generation $0$, as defined in Equation 4 in the main manuscript. As for the hyperparameters in OCS, $t^{\star}$ represents the target generation, $h^{\star}$ indicates the genetic diversity to be maintained in the target generation, and $s$ is the shape parameter that defines the trajectory of $h^{(t)}$. In this study, we set $t^{\star} = T_r = 4$, $h^{\star} = 0.01h^{(0)}$, and $s = 1$.

### Supplemental Figures

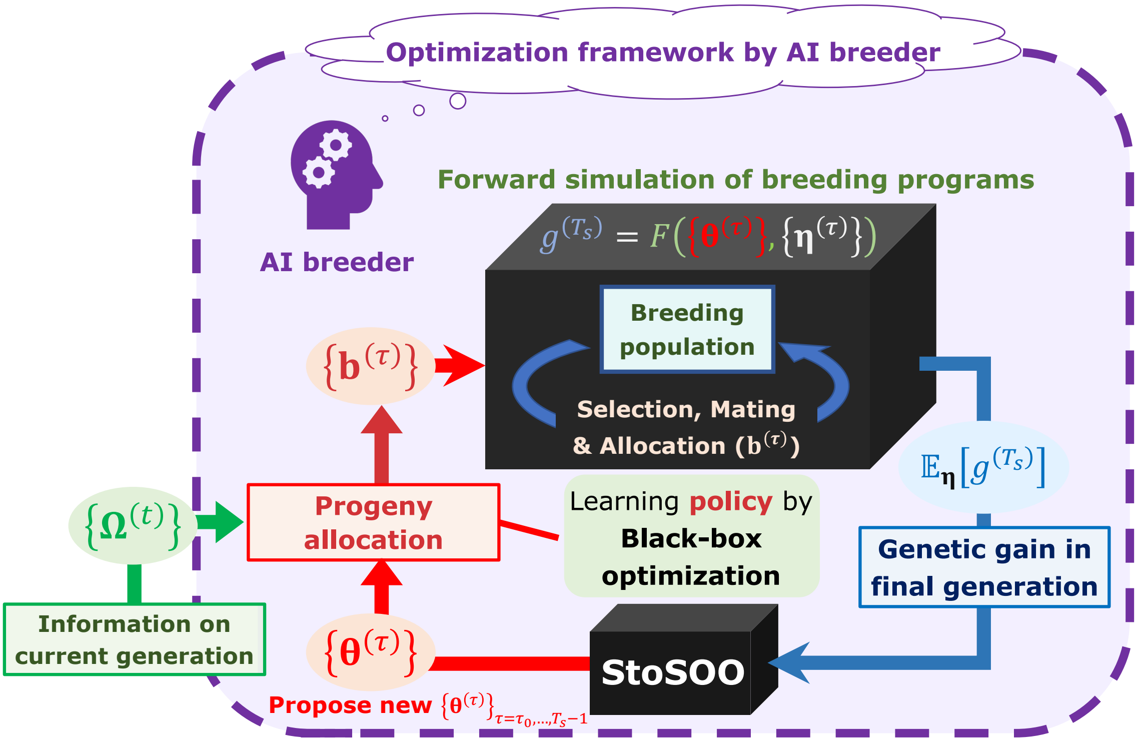


**Supplemental Fig. 1.** Image of the framework to optimize progeny allocation for each mating pair in the breeding scheme. In our framework, the breeders can obtain a set of parameters representing the optimal allocation strategy, $\{\hat{\boldsymbol{\theta}}^{(\tau)}\}_{\tau=\tau_0,\dots,T-1}$, by providing the AI breeder with information on an initial breeding population, as in Hamazaki and Iwata (2024). Here, a breeding simulator is regarded as a function whose input is a set of allocation parameters, and whose output is the final genetic gain. This figure is adapted from the Figure 1 in Hamazaki and Iwata (2024).

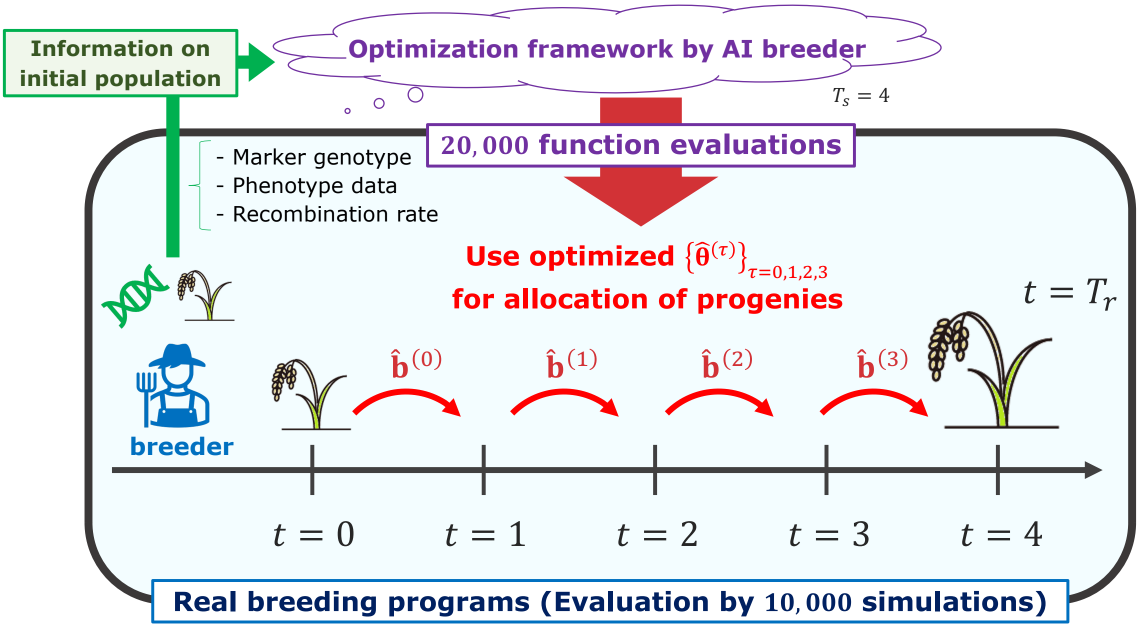


**Supplemental Fig. 2.** Optimization of allocation strategies in breeding schemes without model updates and its evaluation. Adapted from the Figure 1 in Hamazaki and Iwata (2024).

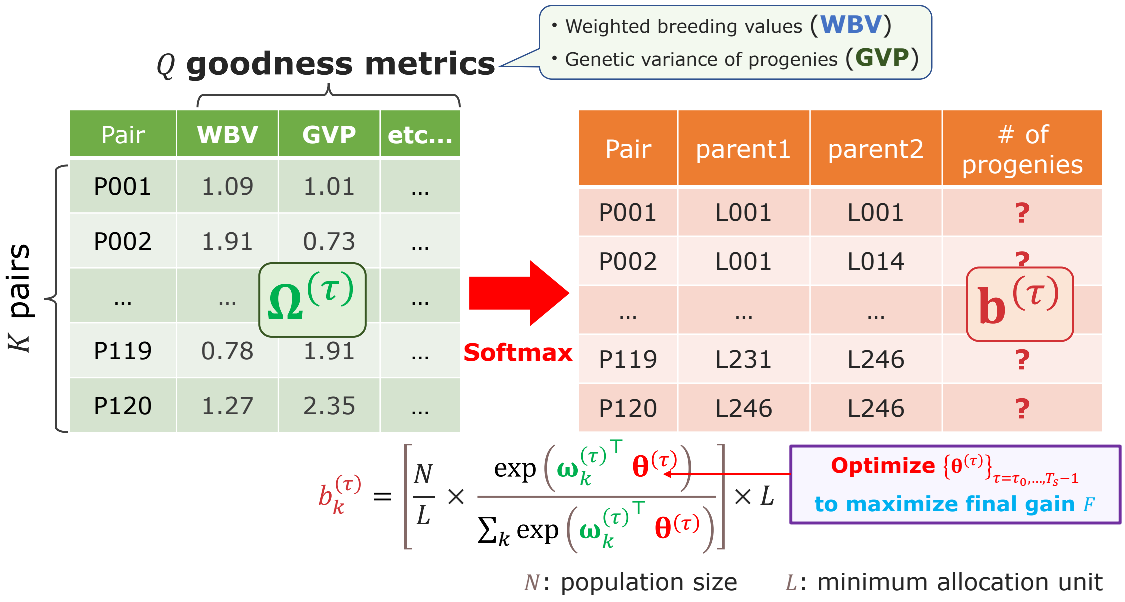


**Supplemental Fig. 3.** The resource allocation strategy of progenies used in this study. After determining mating pairs by the diallel crossing between the parent candidates, given the parameter $\boldsymbol{\theta}^{(\tau)}$, the breeder automatically determines the number of progenies allocated to each pair $b_k^{(\tau)}$. This step is realized by utilizing the matrix $\boldsymbol{\Omega}^{(\tau)}$ consisting of features representing some goodness of mating pairs, i.e., WBV and GVP in this study. In other words, we assumed that $b_k^{(\tau)}$ is determined by applying the softmax function to the weighted sum of the multiple features. This figure is adapted from the Figure 2 in Hamazaki and Iwata (2024).

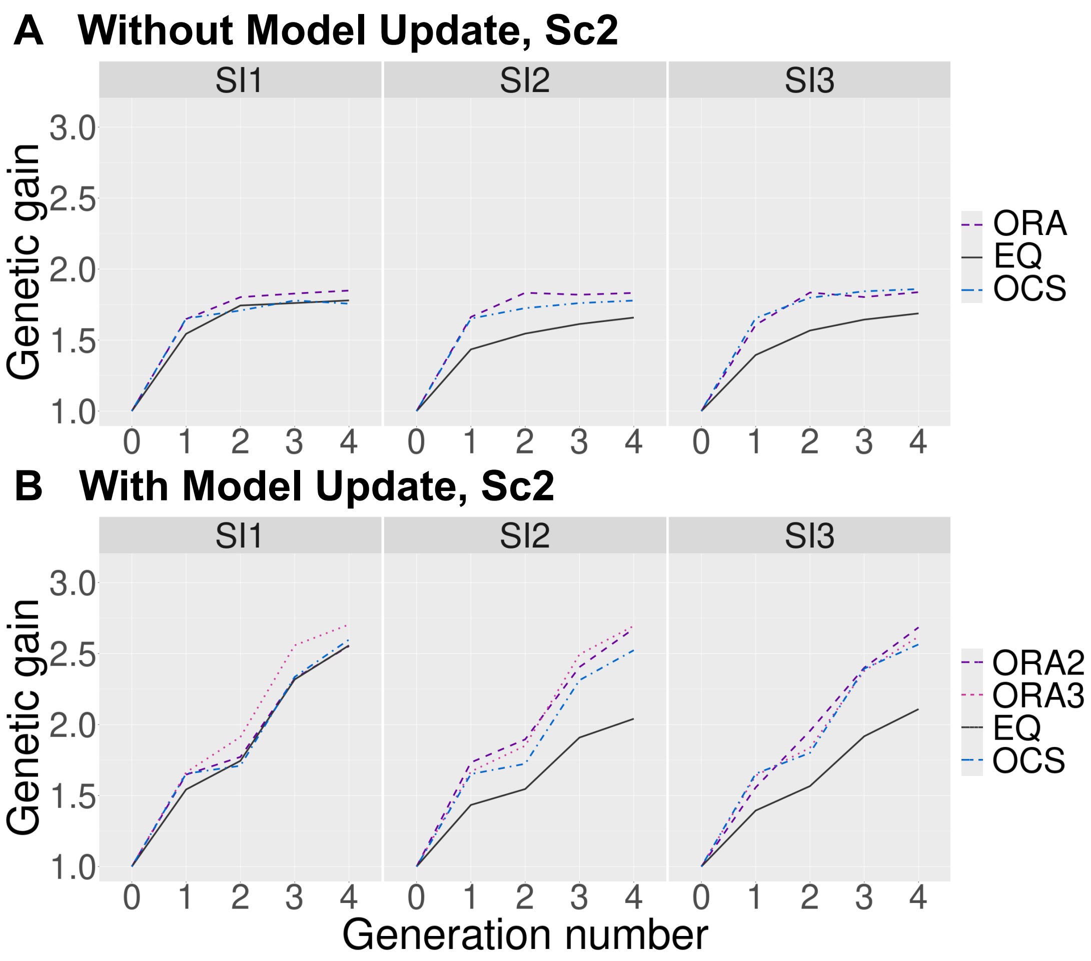


**Supplemental Fig. 4.** Change in the expected genetic gains over four generations under different selection intensities for Scenario 2. The horizontal and vertical axes represent the number of generations and the genetic gains, respectively. Panel (A) corresponds to the scheme without model updates (comparing ORA, EQ, and OCS), while panel (B) corresponds to the scheme with model updates (comparing ORA2, ORA3, EQ, and OCS). The abbreviations of the allocation strategies are the same as those of Fig. 3.

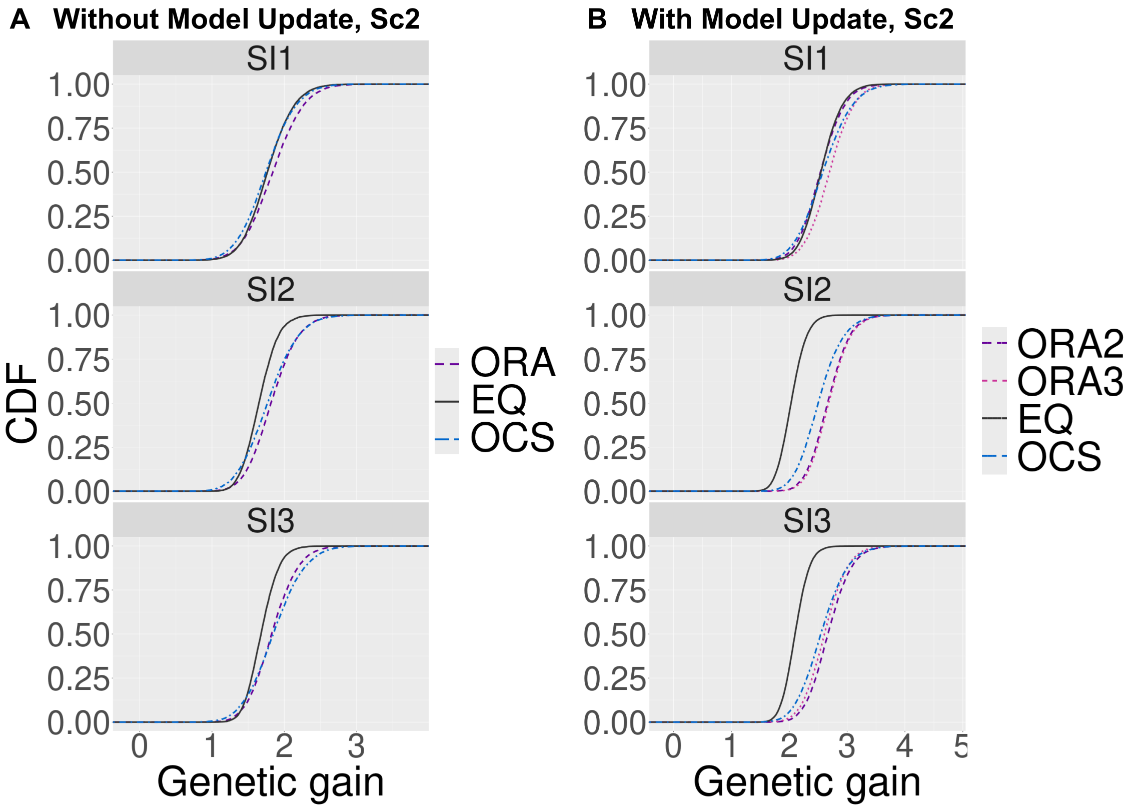


**Supplemental Fig. 5.** Genetic gains across different simulation repetitions for Scenario 2. The horizontal axis shows the final genetic gain of an individual, whereas the vertical axis represents the percentile of the simulation repetitions. Panel (A) corresponds to the scheme without model updates (comparing ORA, EQ, and OCS), while panel (B) corresponds to the scheme with model updates (comparing ORA2, ORA3, EQ, and OCS). The abbreviations of the allocation strategies are the same as those of Fig. 3.

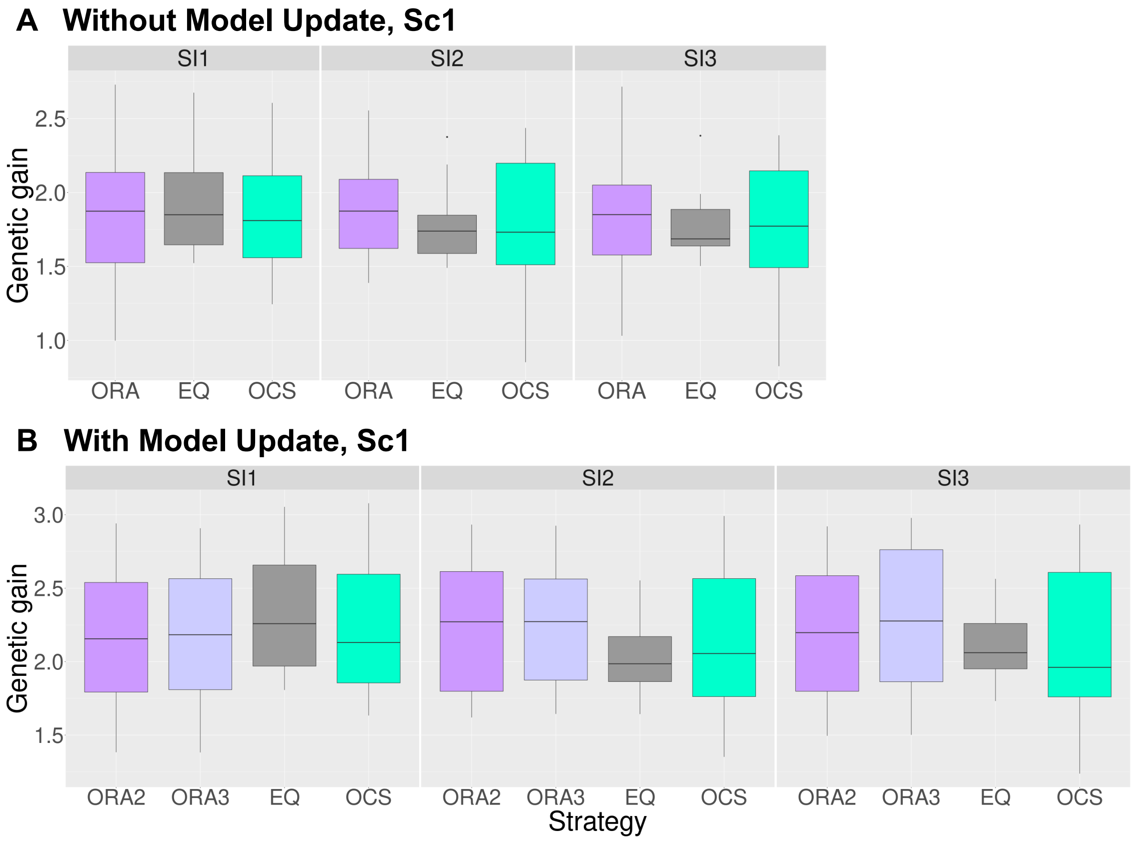


**Supplemental Fig. 6.** Genetic gains in the final generation using ten replications for the phenotype simulation under different selection intensities for Scenario 1. This figure includes boxplots of the genetic gains in the final generation using ten replications for the phenotype simulation. The horizontal and vertical axes represent the different allocation strategies and the final genetic gains, respectively. We compared the following allocation strategies under the different selection intensities (SI1 – SI3): ORA (purple), ORA2 (purple), ORA3 (light purple), EQ (grey), and OCS (light blue), with details given in Fig. 3. Panel (A) corresponds to the scheme without model updates (comparing ORA, EQ, and OCS), while panel (B) corresponds to the scheme with model updates (comparing

ORA2, ORA3, EQ, and OCS).

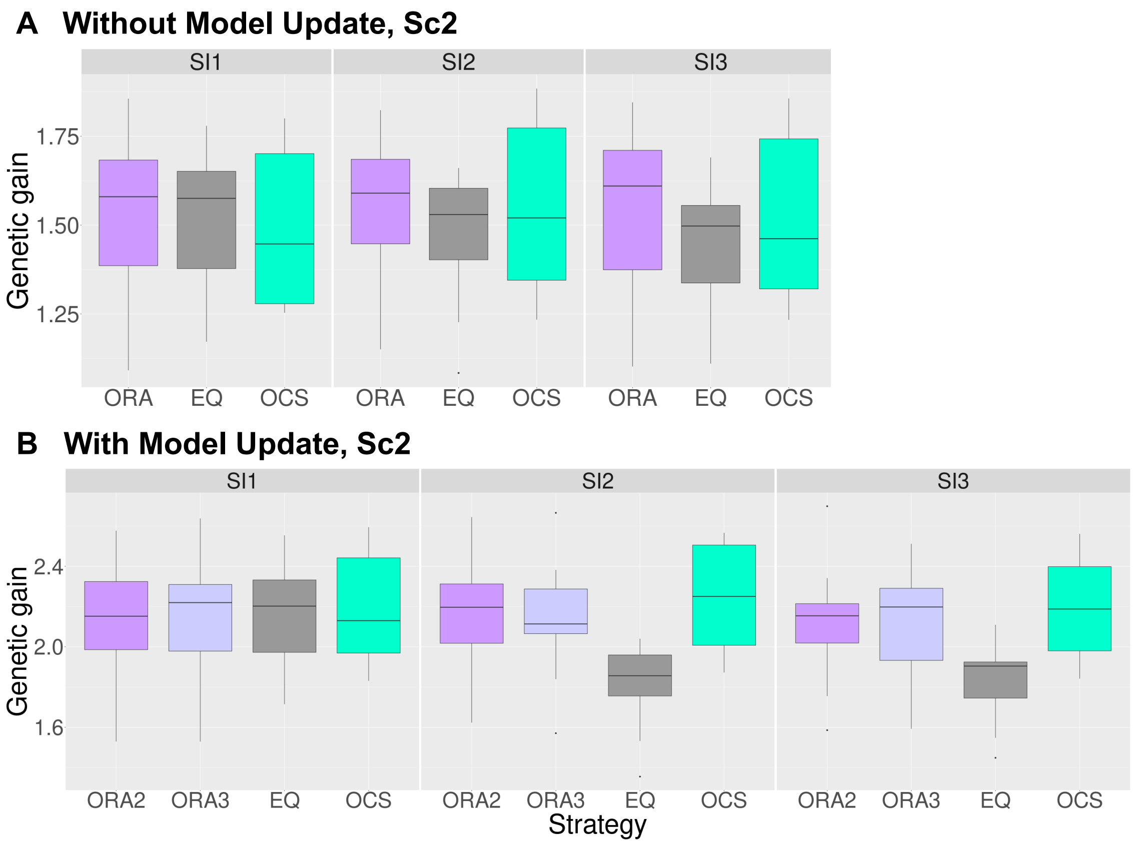


**Supplemental Fig. 7.** Genetic gains in the final generation using ten replications for the phenotype simulation under different selection intensities for Scenario 1. This figure includes boxplots of the genetic gains in the final generation using ten replications for the phenotype simulation. The horizontal and vertical axes represent the different allocation strategies and the final genetic gains, respectively. We compared the following allocation strategies under the different selection intensities (SI1 – SI3): ORA (purple), ORA2 (purple), ORA3 (light purple), EQ (grey), and OCS (light blue), with details given in Fig. 3. Panel (A) corresponds to the scheme without model updates (comparing ORA, EQ, and OCS), while panel (B) corresponds to the scheme with model updates (comparing

ORA2, ORA3, EQ, and OCS).

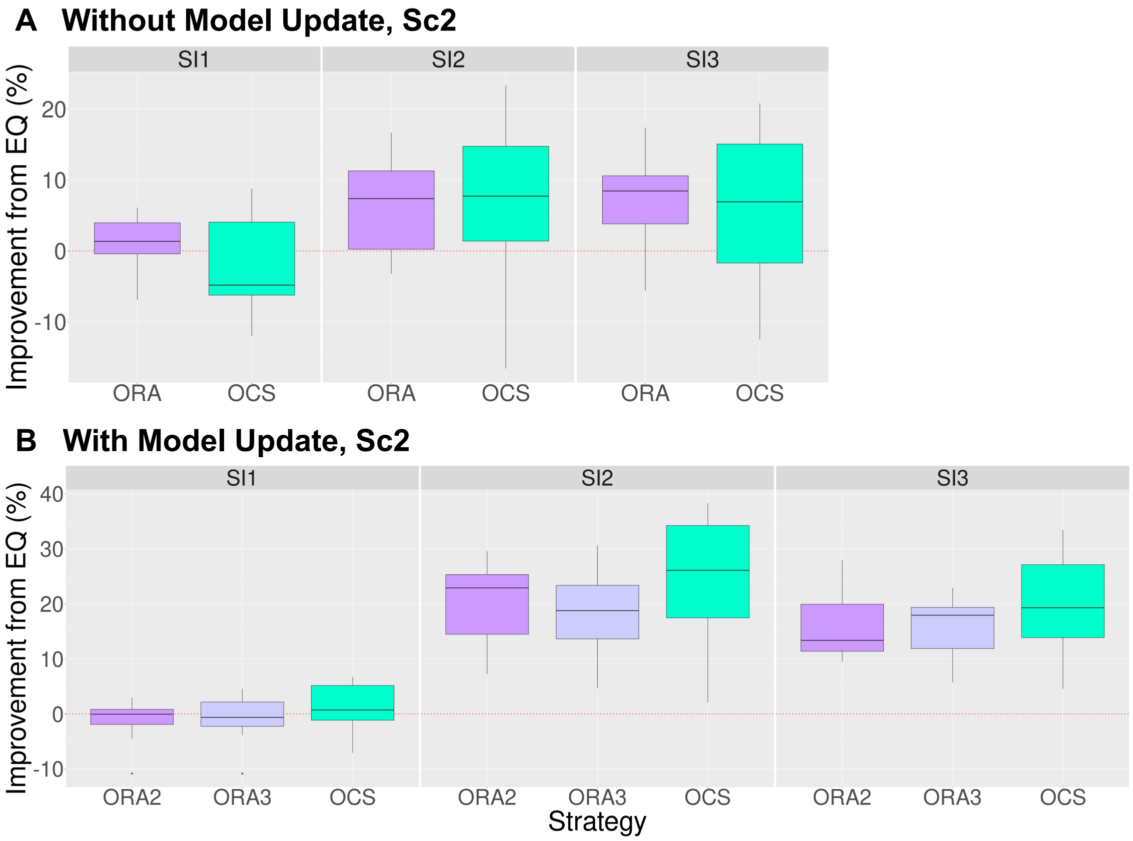


**Supplemental Fig. 8.** Improvement of the final genetic gains compared to the equal allocation strategy using ten replications for the phenotype simulation under different selection intensities for Scenario 2. This figure includes boxplots of the improvement of the final genetic gains compared to the equal allocation strategy using ten replications for the phenotype simulation. The horizontal and vertical axes represent the different allocation strategies and the improvement rate of the final genetic gains, respectively. We compared the following allocation strategies under the different selection intensities (SI1 – SI3): ORA (purple), ORA2 (purple), ORA3 (light purple), and OCS (light blue), with details given in Fig. 3. Panel (A) corresponds to the scheme without model updates (comparing ORA and OCS), while panel (B) corresponds to the scheme with model updates (comparing ORA2, ORA3, and OCS).

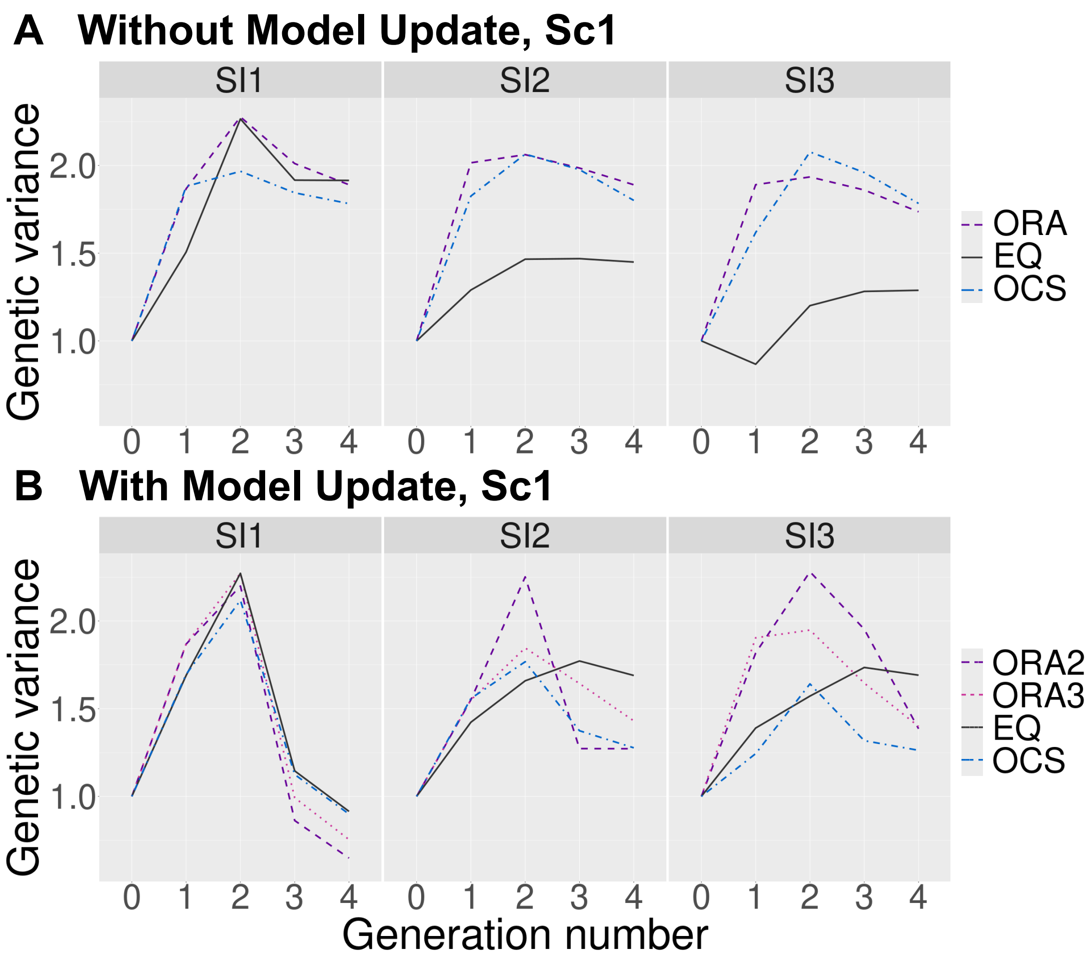


**Supplemental Fig. 9.** Change in the genetic variances over four generations under different selection intensities for Scenario 1. The horizontal and vertical axes represent the number of generations and the genetic variances in the breeding populations, respectively. Panel (A) corresponds to the scheme without model updates (comparing ORA, EQ, and OCS), while panel (B) corresponds to the scheme with model updates (comparing ORA2, ORA3, EQ, and OCS). The abbreviations of the allocation strategies are the same as those of Fig. 3.

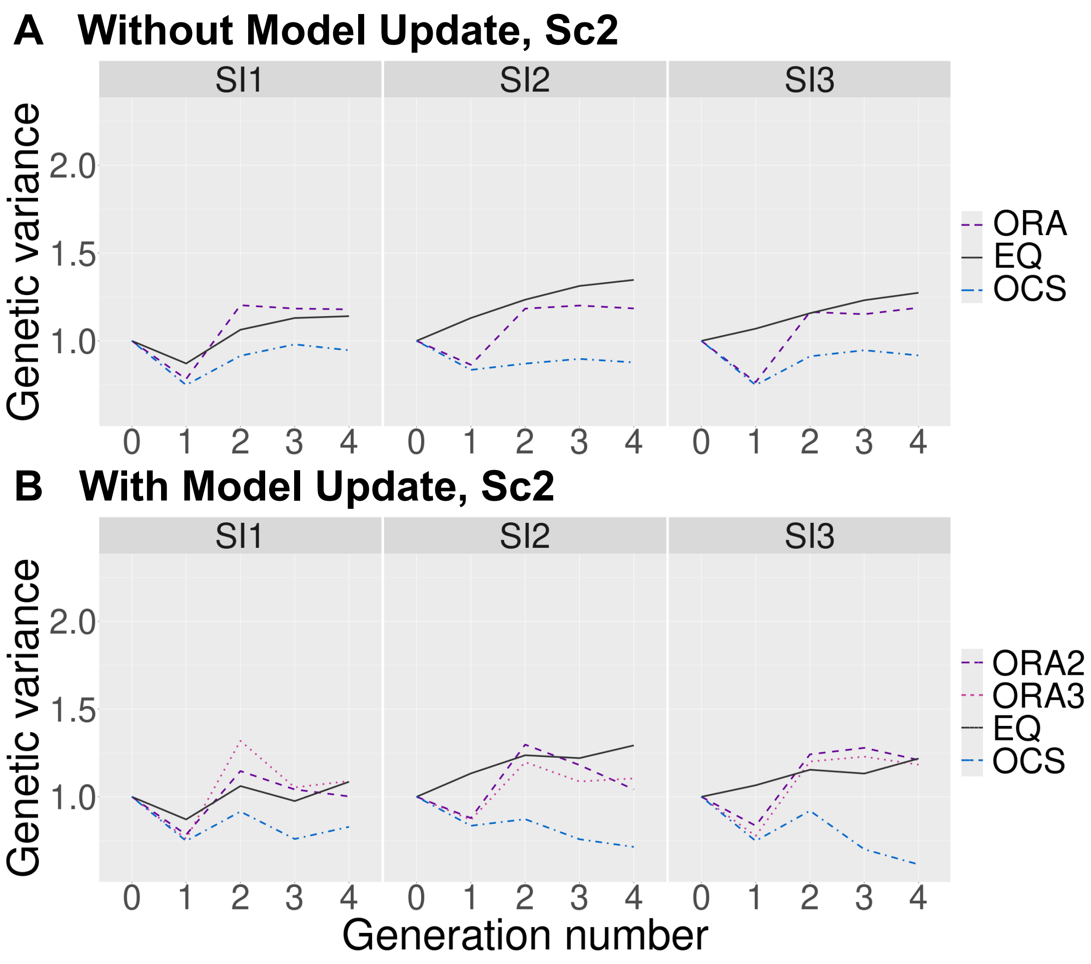


**Supplemental Fig. 10.** Change in the genetic variances over four generations under different selection intensities for Scenario 2. The horizontal and vertical axes represent the number of generations and the genetic variances in the breeding populations, respectively. Panel (A) corresponds to the scheme without model updates (comparing ORA, EQ, and OCS), while panel (B) corresponds to the scheme with model updates (comparing ORA2, ORA3, EQ, and OCS). The abbreviations of the allocation strategies are the same as those of Fig. 3.

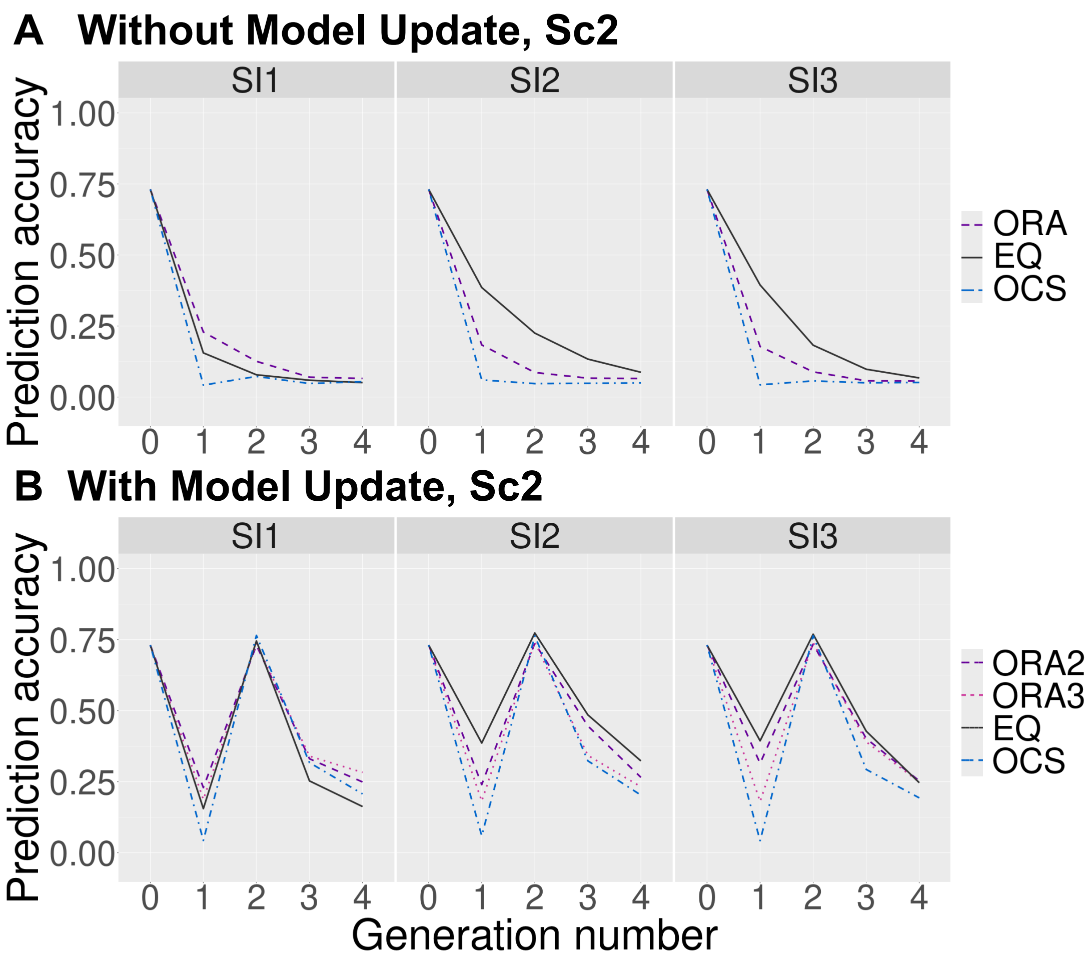


**Supplemental Fig. 11.** Change in the prediction accuracies over four generations under different selection intensities for Scenario 2. The horizontal and vertical axes represent the number of generations and the prediction accuracies for the breeding populations, respectively. Panel (A) corresponds to the scheme without model updates (comparing ORA, EQ, and OCS), while panel (B) corresponds to the scheme with model updates (comparing ORA2, ORA3, EQ, and OCS). The abbreviations of the allocation strategies are the same as those of Fig. 3.
